# Spatial transcriptomic profiling of the human aortic valve reveals cellular sex differences near sites of calcification

**DOI:** 10.1101/2025.08.20.671175

**Authors:** Talia Baddour, Van K. Ninh, Rayyan M. Gorashi, Rene Peña, Kevin R. King, Brian A. Aguado

## Abstract

Sex differences in aortic valve stenosis (AVS) progression have been documented clinically, but the underlying cellular mechanisms that drive sex-dependent calcification in aortic valve tissue remain poorly understood. Here, we harnessed single cell and spatial transcriptomics to investigate mechanisms that drive sex dependent spatial organization of valvular interstitial cell (VIC) and macrophage gene expression near calcification sites in human male and female aortic valve tissue. Histological analyses of aortic valve tissues stratified into healthy and diseased cohorts based on degree of calcification reveal increased valve calcification area in diseased male aortic valves relative to female, and increased valve thickening in diseased female aortic valves. Single cell sequencing analysis of heterogeneous valvular interstitial cell (VIC) populations reveals male-dependent gene expression of the Activator Protein 1 (AP-1) transcription factor complex. Spatial transcriptomics and RNA-FISH analyses of VIC populations near sites of calcification revealed male-dependent gene expression localization of Cartilage Oligomeric Matrix Protein (*COMP*), as opposed to diffuse *COMP* expression in female VICs. Cell-cell communication analyses were used to determine female-specific macrophage-VIC interactions. Secreted phosphoprotein 1 (also known as osteopontin) expressed from macrophages interacts with the cell surface receptor CD44 expressed by VICs to drive a pro-fibrotic phenotype in female aortic valves. Together, our results reveal sex differences in VIC and macrophage heterogeneity and functions near sites of calcification in aortic valve tissue. Our results highlight the importance of sex-based transcriptomics analyses to understand the cellular phenotypes responsible for causing sex differences in aortic valve fibrosis calcification.

## Introduction

Aortic valve stenosis (AVS) is a progressive and life-threatening condition characterized by narrowing and stiffening of the aortic valve due to fibro-calcification^1–3^. AVS is also a sex-dependent disease, where numerous clinical and imaging studies have reported sex differences in progression and outcomes to valve replacements. For instance, male AVS patients typically exhibit greater ectopic calcification and more rapid deterioration of the aortic valve, whereas female patients with equivalent disease severity present with more fibrotic valve thickening and preserved aortic valve area^4–6^. Sex differences in aortic valve tissue persist even after adjusting for age and comorbidities, suggesting intrinsic biological mechanisms that contribute to modulating sex differences in AVS progression. Recognizing that female patients are less frequently recommended for aortic valve replacement even after adjusting for increased AVS incidence in male patients^7^, there is a critical need to determine the sex-specific mechanisms of AVS progression and identify alternative paths for non-surgical interventions.

Sex-specific differences in valvular interstitial cells (VICs) have been consistently reported in human tissues and animal models of aortic valve stenosis^8–12^. As such, VICs likely play a central role in modulating sex-dependent extracellular matrix composition, remodeling, and fibro-calcification of the aortic valve. VICs are the primary stromal cell of the aortic valve and are responsible for maintaining valve homeostasis under healthy physiological conditions^13–16^. VICs also activate to pathogenic myofibroblasts and can further differentiate into osteoblast-like cells in response to mechanical stress or inflammation^17^. Histological analyses have revealed that male aortic valves tend to exhibit higher levels of calcium phosphate accumulation^18^ and expression of Y-linked demethylases^9^, whereas female valves have increased collagen abundance and myofibroblast activation likely due to increased Rho/ROCK signaling via X-linked gene expression^8^, estradiol receptor activity^19,20^ and immune-VIC interactions^21–23^. Despite our current understanding of sex differences in VIC phenotypes, there remains a gap in understanding how male and female VICs are organized within the aortic valve to modulate sex-dependent tissue structures during AVS progression.

Myofibroblast heterogeneity is vast, and a challenge that persists is understanding how biological sex contributes to heterogenous cell-cell and cell-matrix interactions in the aortic valve. Aortic valve tissue is organized into a tri-layer structure, where the fibrosa, spongiosa, and ventricularis contain a mixture of cell types including VICs, valvular endothelial cells (VECs), immune cells, and other resident stromal cell populations^24,25^. Recent advances in single cell and histological analyses have revealed heterogeneity in valve cell populations and their spatial organization within the aortic valve^1,26^. However, the categorization of myofibroblasts by upregulated genes or surface markers alone may not capture the functional activity of myofibroblasts or osteoblast-like cells within tissues. One common characteristic that pathogenic myofibroblasts have in common is their spatial proximity to key features of aortic valve tissue, namely calcium phosphate particles^27^, fibrillar collagen^24^, inflammatory immune cells^28^, and damaged endothelium^29^. We posit myofibroblasts and osteoblast-like VICs that are near active sites of calcification are likely the main pathogenic cells that drive AVS progression, prompting us to understand sex-dependent valve cell heterogeneity as a function of spatial location in the aortic valve.

Here, we leveraged single cell and spatial transcriptomics to generate sex-specific cell atlases from human aortic valves and identified male and female VIC populations near sites of calcification. We obtained healthy and calcified aortic valve tissue from male and female human donors and systematically characterized sex-dependent spatial gene expression patterns in histologically defined tissue regions. First, we confirmed male aortic valve leaflets have increased calcification relative to female leaflets which had increased leaflet thickness. Next, we generated single cell and spatial transcriptomics datasets to reveal sex-biased VIC heterogeneities in aortic valve tissue, in addition to their spatial organization of myofibroblast and osteoblast-associated genes in diseased aortic valves. Specifically, we identified and validated expression of cartilage oligomeric matrix protein (*COMP*) localized to sites of calcification uniquely in male VICs. In female valves, we identified macrophages expressing *SPP1* interact with the glycoprotein receptor CD44 on VICs to drive fibrotic remodeling in female valve tissues. Together, our spatial transcriptomics analyses of the human aortic valve have provided insights into the spatial organization of pathogenic cell types that modulate sex-dependent aortic valve calcification.

## Results

### Histological analysis of human aortic valve tissue reveals sex differences in valve calcification and morphology

We analyzed human aortic valve tissue samples via histology to corroborate clinical findings demonstrating sex differences in aortic valve fibro-calcification^4,6,18^. We collected valves from female (N = 9) and male (N = 12) patients with varying risk factors for disease (**Supplementary Table I**). We utilized Alizarin Red and Masson’s Trichrome reagents to stain cryosectioned aortic valve leaflets (**Figure 1A – 1B, Supplementary Figure I**) and stratified healthy and diseased patients based on Warren-Yong scoring (1: absent, 2: mild valve thickening and early nodular calcification, 3: moderate thickening with many calcified nodules, 4: severe thickening with many calcified nodules) as previously described^31^ (**Figure 1C, Supplementary Figure II).** We observed heterogenous calcium surface area and dense fiber presence among male and female patients and no significant correlation between dense fibers and calcium surface area for male tissues (R^2^ = 0.304, *P* = 0.0783) or female tissues (R^2^ = 0.000837, *P* = 0.946) (**Supplementary Figure III**). We confirmed no significant differences in age or disease risk factors between sex separated diseased female and diseased male patient cohorts (**Supplementary Table IX**).

**Figure 1.**
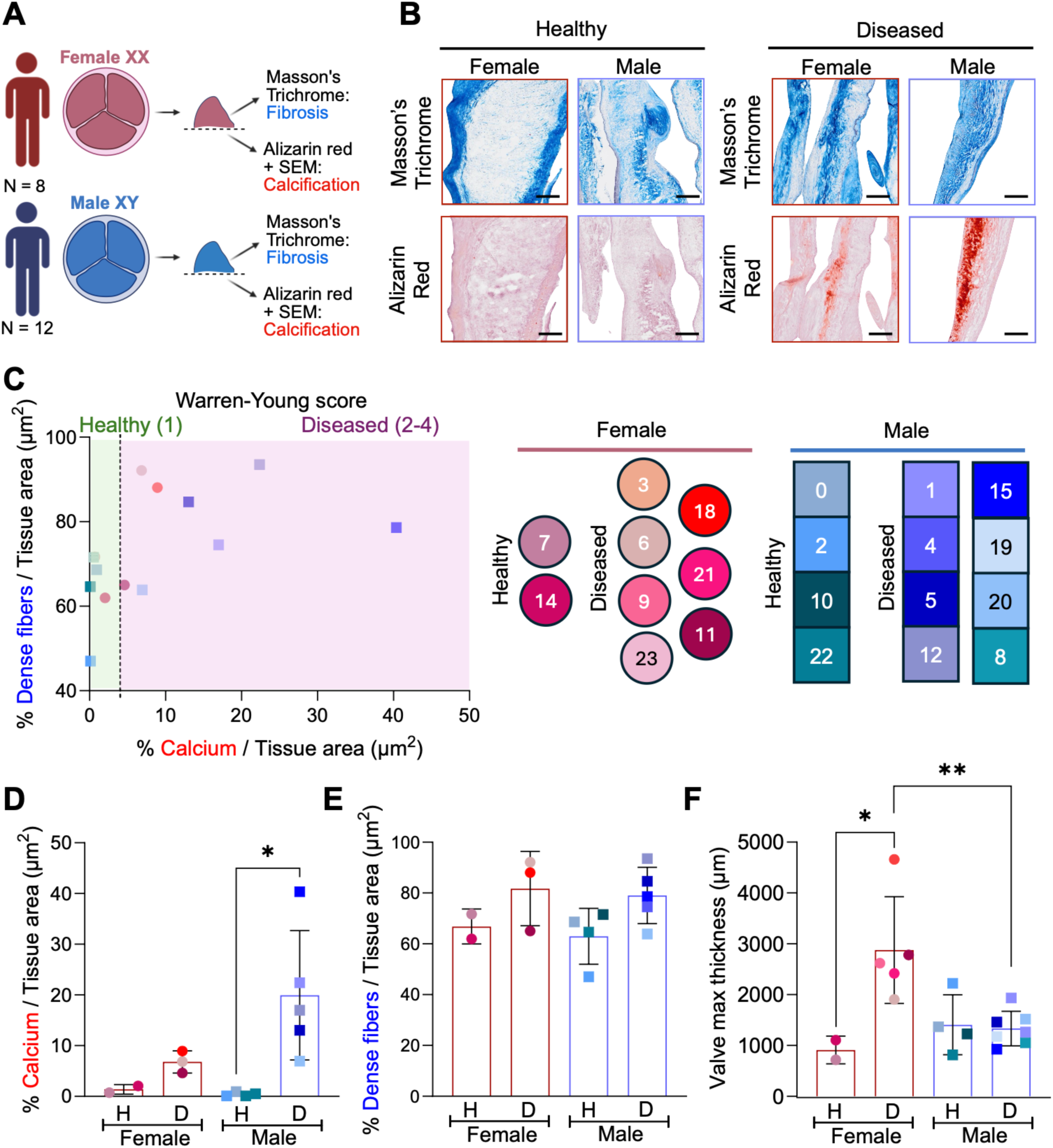
Histological analysis of sex differences in fibro-calcification of human aortic valves. **A.** Schematic of experimental design for histological characterization of male (N = 12) and female (N = 9) human aortic valves. **B.** Representative images of healthy and diseased aortic valve cross sections stained with Masson’s trichrome to characterize collagen fiber density with Alizarin red to characterize calcification. Scale bar = 100 μm. **C.** Stratification of patients into healthy and diseased cohorts based on Warren-Yong scoring (shape color and number indicate different patients described in **Supplementary Table I**). **D.** Percent calcium over tissue area for male (N = 9) and female (N = 5), healthy (H, N = 6) and diseased (D, N = 8) patients. Means ± standard deviations and *P*-values of comparisons shown with statistical significance determined via two-way ANOVA with Tukey posttests. \**P* < 0.05. **E.** Percent dense fibers over tissue area for male (N = 9) and female (N = 5), healthy (H, N = 6) and diseased (D, N = 8) patients. Means ± standard deviations and *P*-values of comparisons shown with statistical significance determined via two-way ANOVA with Tukey posttests. **F.** Maximum aortic valve thickness measurements for each patient. Means ± standard deviations and *P*-values of comparisons shown with statistical significance determined via two-way ANOVA with Tukey posttests. \**P* < 0.05. \*\**P* < 0.01.

After categorizing patients as healthy or diseased, we hypothesized measures of calcification to be increased in male patients and measures of fibrosis to be increased in female patients. We quantified sex differences in calcium area, collagen fiber density, and valve thickness to test our hypothesis. Calcium surface area quantification revealed a male-specific significant increase in percent calcified tissue area in diseased male tissue (19.9 ± 12.7%) relative to healthy male tissue (0.4 ± 0.4%) (**Figure 1D**). Diseased female aortic valve tissue had trends toward increased calcified tissue area (6.8 ± 2.2%) relative to healthy specimens (1.4 ± 0.9%). Further analysis showed that male valves with Warren-Yong score of 3 have increased calcium surface area (31.4 ± 12.7%) relative female valves in the same category (7.9 ± 1.5%). Even though no significant differences were observed in fiber density (**Figure 1E**), diseased female aortic valve tissue had a sex-specific increase in maximum valve thickness (2877 ± 1050 μm) relative to diseased male aortic valves (1333 ± 337.7 μm) (**Figure 1F**). When comparing within sex, we observed a female-dependent increase in valve thickness in female diseased tissue relative to healthy (911.5 ± 273.7 μm), whereas diseased male and healthy male tissues (1408 ± 591.1 μm) had no significant differences in valve thickness (**Figure 1F**). We also note the ratio of percent calcification relative to tissue thickness was increased uniquely in the male diseased cohort (**Supplementary Figure IV)**.

We also performed scanning electron microscopy to quantify sex differences in calcium phosphate particle morphology and size distributions (**Supplementary Figures VA – VD).** Average particle diameters revealed no sex differences between diseased female valves (mean = 515 ± 57.3 nm) and diseased male valves (mean = 397 ± 178 nm) (**Supplementary Figure VD).** However, the range in particle size for male tissues was ∼2-fold wider (379.4 nm) compared to female particle size (155.4 nm), revealing greater particle size heterogeneity in male valves relative to female. Taken together, we identified sex differences in tissue calcification and morphology which led us to investigate the cell phenotypes that may cause the observed differences.

### Single cell RNA sequencing and spatial transcriptomics reveal sex differences in valve cell heterogeneity driving valve fibro-calcification

We sought to define sex differences in valve cells near sites of calcification using single cell and spatial transcriptomics, given our observations of increased calcification burden in males relative to females. We integrated our own single cell RNA sequencing data with publicly available data sets from a total of N = 4 female and N = 8 male patients (**Figure 2A**). Uniform Manifold Approximation and Projection (UMAP) plots of cells from integrated single cell sequencing data shows uniform integration across 61,853 patient cells (**Figure 2B**). We next performed a pseudo-bulk RNA seq analysis on healthy vs. diseased valve cells and compared significant genes in our dataset to genes that encode for proteins in previously published proteomic datasets defining fibrosis and calcification in diseased aortic valve tissue^1^ (**Figures 2C – 2D).** We observed sex-specific overlap of genes associated with fibrosis-related proteins, including *LGALS1, MIF, PLA2G2A, SERPINE2,* and *S100A11* for male valves and *ACTB, COL3A1, TAGLN2, CCL18, S100A10* for female valves. We also observed sex differences in gene overlap with calcification proteins such as *TIMP1, HLA-A, MYL6, TIMP1, IGFBP4* in male valves and *FN1, C1QB, CST3, SPARC,* and *HLA-DP1* for female valves. A complete list of overlapping genes is available in **Supplementary Table X**.

**Figure 2.**
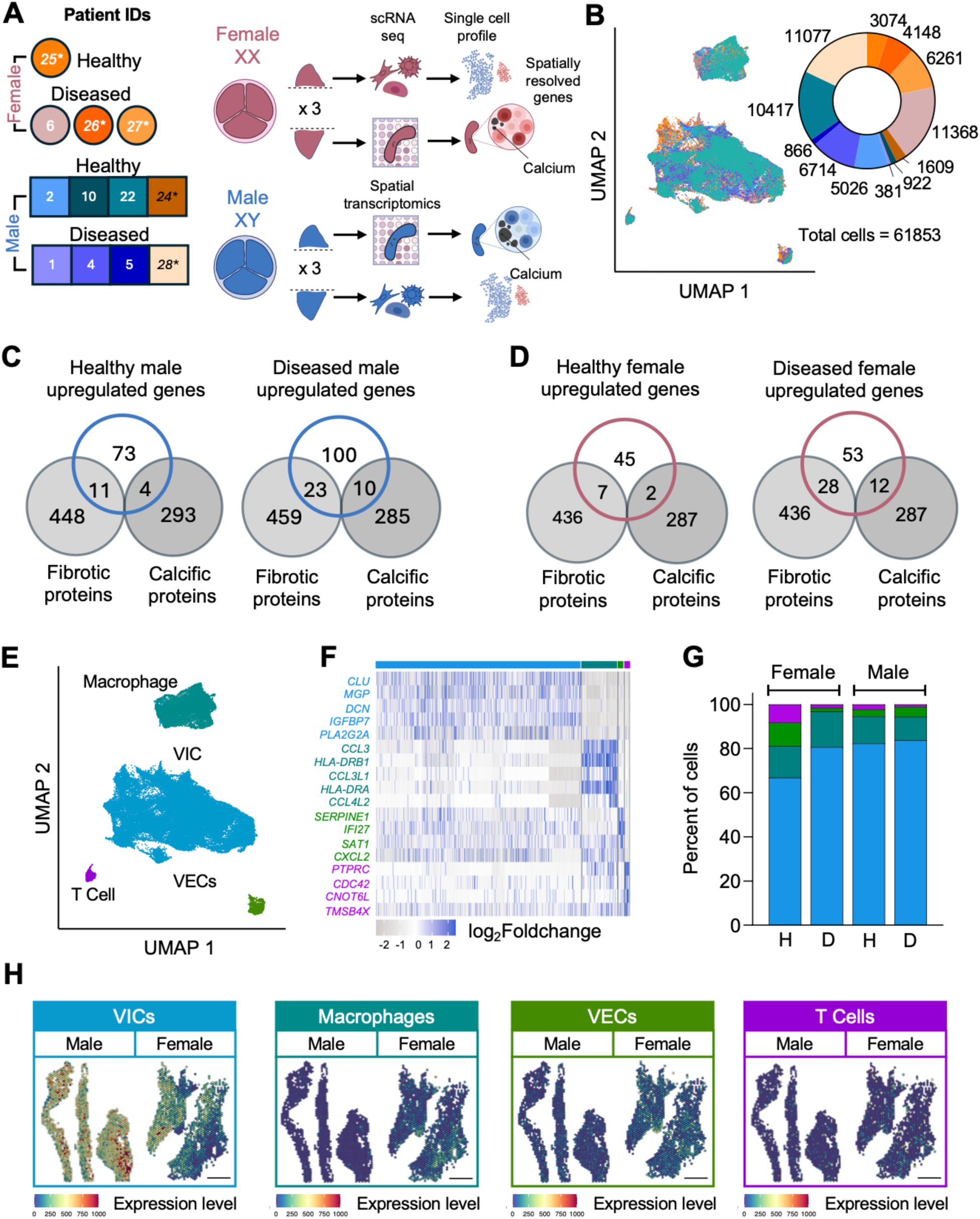
Single cell RNA sequencing and spatial transcriptomics analysis of human aortic valves. **A.** Schematic describing experimental workflow for patient valves used for single cell RNA sequencing and spatial transcriptomics analyses; female healthy (N = 1), female diseased (N = 3), male healthy (N = 4), male diseased (N = 4). Asterisks indicate single cell sequencing data obtained from Xu et. al. 2020^26^. **B.** Unsupervised clustering of 61853 aortic valve cells. Pie chart describes distribution of cell populations by patient ID color. **C.** Venn diagrams comparing upregulated genes in male healthy (N = 4) and diseased (N = 4) aortic valves identified via pseudo-bulk RNA sequencing analysis to catalogs of 459 fibrosis associated and 297 calcification associated proteins identified from Schlotter et. al. 2018^1^. **D.** Venn diagrams comparing upregulated genes in female healthy (N = 1) and diseased (N = 3) aortic valves identified via pseudo-bulk RNA sequencing analysis to catalogs of 459 fibrosis associated and 297 calcification associated proteins identified from **E.** Unsupervised clustering of aortic valve cells identified by cell type. **F.** Top 5 gene markers per cell cluster. Significance determined using Wilcoxon rank sum testing; log_2_Foldchange > |0.5|; \**P*_adj_<0.0001. **G.** Proportions of cell types across combined patient groups: female healthy (N = 1), female diseased (N = 3), male healthy (N = 4), male diseased (N = 4). Shaded colors in bars correspond to colors used to label cells in unsupervised clustering plot. **H.** Top 5 genes per cell cluster scored onto spatial transcriptomics map of female diseased and male diseased tissue. Scale bar = 1000 μm.

We next determined aortic valve cell populations and their spatial location within male and female aortic valve leaflets. Unsupervised clustering of cells isolated from aortic valve leaflets revealed 4 distinct populations valve cells and their associated biomarkers: [1] valvular interstitial cells (VICs) (*CLU, MGP, DCN, IGFBP7, PLA2G2A*), [2] macrophages (*CCL3, HLA-DRB1, CCL3L1, HLA-DRA, CCL4L2*), [3] valvular endothelial cells (VECs) (*SERPINE1, IFI27, SAT1, CXCL2*), and [4] T cells (*PTPRC, CDC42, CNOT6L, TMSB4X*) (**Figures 2E – 2F**). We observed diseased cell populations were similar between sex (**Figure 2G**). Next, our spatial transcriptomics analyses of aortic valve tissue enabled us to spatially map cell populations on sections of aortic valve tissue. Valve scoring of the top 5 genes for valvular VICs, macrophages, VECs, and T cells demonstrates valve specific cellular heterogeneity across diseased valves (**Figure 2H**). In sum, our multi-omics analyses identified sex-dependent expression of genes associated with fibro-calcification and the spatial organization of cell populations in aortic valves.

### VIC sub-clustering analysis reveals male-specific VIC association with the transcription factor Activator Protein 1

We next investigated sex differences in VIC transcriptomic heterogeneity, since VICs represent the largest population in our single cell sequencing datasets that likely modulate sex-dependent fibro-calcification. To probe fibroblast heterogeneity in diseased valve tissue, we subclustered the VIC population into 4 subtypes via marker expression (**Figure 3A**): VIC 1 (*COLA1A1, COL3A1, COL1A2, COMP, SPARC*), VIC 2 (*PLA2G2A, MT1X, CHI3L2, MT1M, HILPDA*), VIC 3 (*G0S2, PTGDS, CXCL1, CFD, TNFAIP6*), and VIC 4 (*CCL2, CCL7, CXCL2, CXCL3, SOD2*) (**Supplementary Figure VI**). We observed VIC subtype 1 is increased in female diseased valves relative to male diseased valves (χ^2^ = 14.99, 1 df, *P* = 0.0001). Additionally, VIC 1 makes up the highest proportion of cells in the valve relative to other VIC subclusters irrespective of sex (**Figure 3B**). VIC 1 has significant gene ontology (GO) terms associated with extracellular matrix organization and bone development (**Figure 3C**). Within the diseased populations, we observe a higher proportion of VIC 2 cells in males relative to females via chi-squared (χ^2^) tests (χ^2^ = 21.31, 1 df, *P* < 0.0001), relative to the higher proportion of VIC 4 cells in females (χ^2^ = 17.38, 1 df, *P* < 0.0001). Significant GO (*P* < 0.0001) terms associated with VIC 2 include cellular response to zinc ion and stress response to metal ion whereas VIC 3 is associated with positive regulation of cell adhesion and leukocyte cell-cell adhesion. Within the male VIC populations, we observe a higher proportion of VIC 1 and VIC 2 cells in healthy males (χ^2^ = 113.0, 1 df, *P* < 0.0001), (χ^2^ = 81.65, *P* < 0.0001) and a higher proportion of VIC 4 cells in diseased males (χ^2^ = 20.67, 1 df, *P* < 0.0001). Within the female populations we observe a higher proportion of VIC 3 in healthy females relative to diseased females (χ^2^ = 26.63, 1 df, *P* < 0.0001). Significant GO terms associated with VIC 3 include neutrophil migration and cell chemotaxis. From our analysis, we identified 4 VIC subtypes associated with ECM remodeling, response to oxidative stress, chemokine expression, and immune cell interactions.

**Figure 3.**
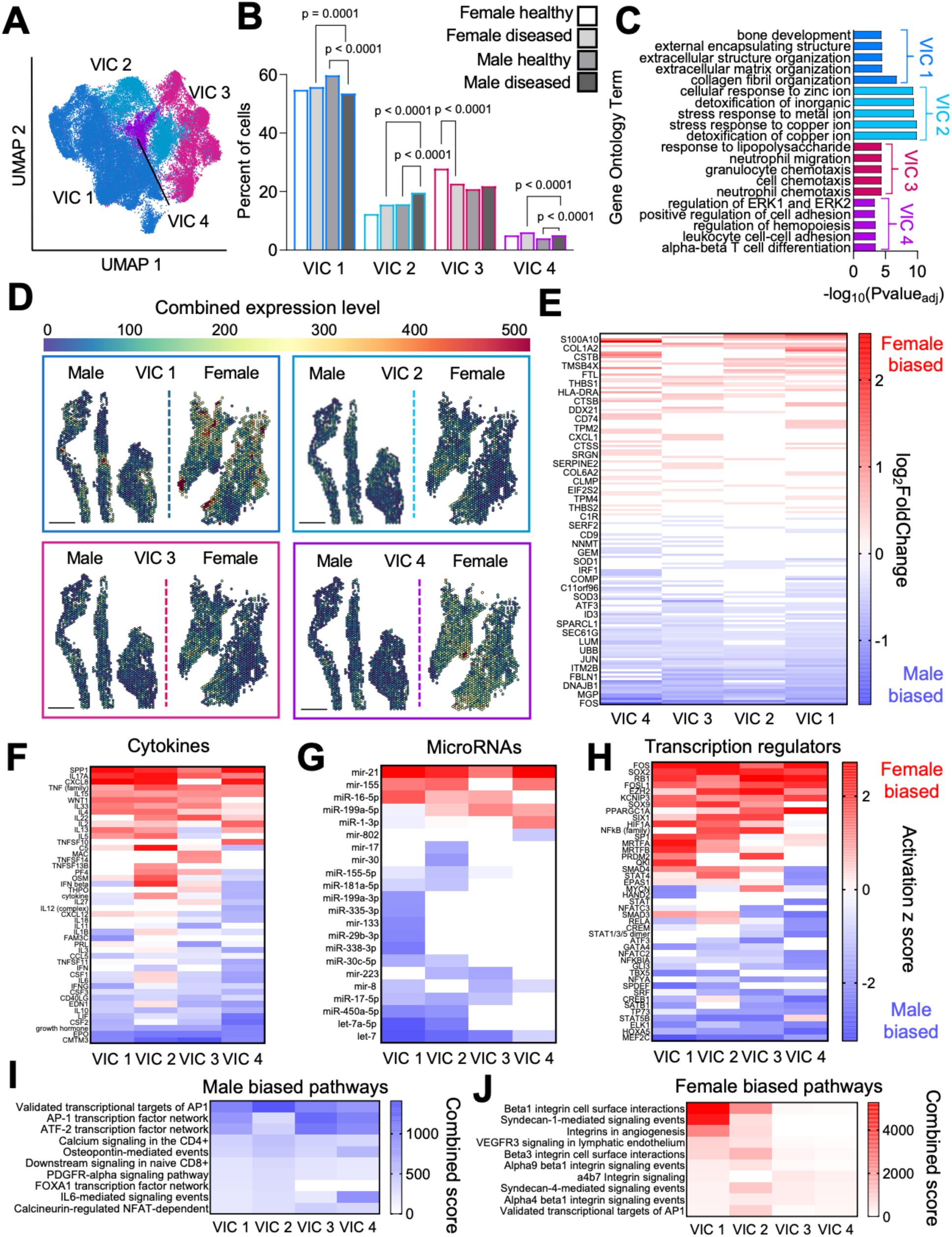
Single cell sequencing reveals sex differences in valvular interstitial cells. **A.** UMAP of subclustered VIC populations. **B.** Proportions of VIC subpopulations for female healthy (N = 1), female diseased (N = 3), male healthy (N = 4), male diseased (N = 4). *P*-values of comparisons shown with statistical significance determined via chi-squared analysis. **C.** Top 5 Gene ontology terms for each VIC subtype. **D.** Top 5 single cell RNA sequencing gene marker for each VIC subtype scored onto spatial transcriptomics maps. Scale bar = 1000 μm. **E.** Differentially expressed genes across all VIC subtypes. Significance determined using Wilcoxon rank sum testing; log_2_Foldchange > |0.5|; \**P*_adj_<0.0001. **F.** Ingenuity Pathway Analysis of cytokines upstream of differentially expressed genes; \**P*_adj_<0.05; activation z-score > |1|. **G.** Ingenuity Pathway Analysis of microRNAs upstream of differentially expressed genes; \**P*_adj_<0.0001; activation z-score > |1|. **H.** Ingenuity Pathway Analysis of transcription regulators upstream of differentially expressed genes; \**P*_adj_<0.05; activation z-score > |1|. **I.** Top 10 EnrichR pathways associated with male biased genes; log_2_Foldchange > |0.5|; \**P*_adj_<0.0001. **J.** Top 10 EnrichR pathways associated with female biased genes; log_2_Foldchange > |0.5|; \**P*_adj_<0.0001.

We next determined where VIC subclusters were located within diseased aortic valve tissue and investigated gene expression differences in VIC subclusters. We scored the top 5 VIC markers for each subcluster onto spatial transcriptomics maps and observed heterogenous localization of VIC subclusters, with VIC 1 mapping to both male and female tissues and VIC 2 and VIC 4 mapping more to female tissue relative to male (**Figure 3D**). Differential gene expression analysis of each VIC subcluster revealed male VICs robustly express *FOS* and *JUN,* which form the Activator protein 1 (AP-1) transcription factor complex known to play a role in osteoblast-like differentiation^36^ (Log_2_FoldChange > |0.5|, *P*_adj_<0.0001) (**Figure 3E**). We also utilized Ingenuity Pathway Analysis (IPA) to reveal sex-biased upstream regulators of VIC subtypes from our differential gene expression analysis. We observed female-biased cytokine-coding genes *SPP1* and *IL17A*, and male-biased cytokine-coding genes *CMTM3* and *EPO* are upstream of genes expressed by all VIC subpopulations (**Figure 3F**). Additionally, we observed an increase in male-biased microRNA regulators uniquely in VIC 1 compared to other VIC subtypes (**Figure 3G**). We also observed female-biased and male-biased expression of transcription factor regulators in all VIC subpopulations (**Figure 3H**). EnrichR pathway analysis confirmed male VICs are biased towards AP-1 transcription factor signaling while female VICs participate in increased integrin cell surface interactions (**Figure 3I – 3J**). Together, our focused analysis on heterogeneous VIC populations revealed sex-dependent pathways and associated gene sets that may differentially modulate fibrosis and calcification.

### VIC 1 marker COMP gene clusters near sites of calcification in male tissues

As a next step, we provided insights into the spatial organization of VICs within the aortic valve and their proximity to regions of calcification. We utilized spatial transcriptomics to reveal gene sets near sites of calcification. First, serial sections of aortic valve tissue reveal cell nuclei near regions of calcification in male and female tissue (**Figure 4A**). We utilized an unsupervised method, Moran’s *I* test statistic, to evaluate highly clustered genes within the tissue. We observed the *COMP* gene ranks as the most clustered gene in male tissue (*I* = 0.39) whereas it ranks as the 9^th^ most clustered gene in female tissue (*I* = 0.45) (**Figure 4B**). We also observed the following 4 top ranking genes in male tissue (*GFAP, PTGDS, GEMIN6, CCN1)* fall in the 9569^th^, 565^th^, 9317^th^, and 218^th^ ranks in the female tissue. We observe the same trends in female tissue with the top 5 ranking genes (*CXCL8, MALAT1, PI3, CLU, CXCL5*) ranking as the 13667^th^, 11^th^, 15^th^, 109^th^ categories in male tissue.

**Figure 4.**
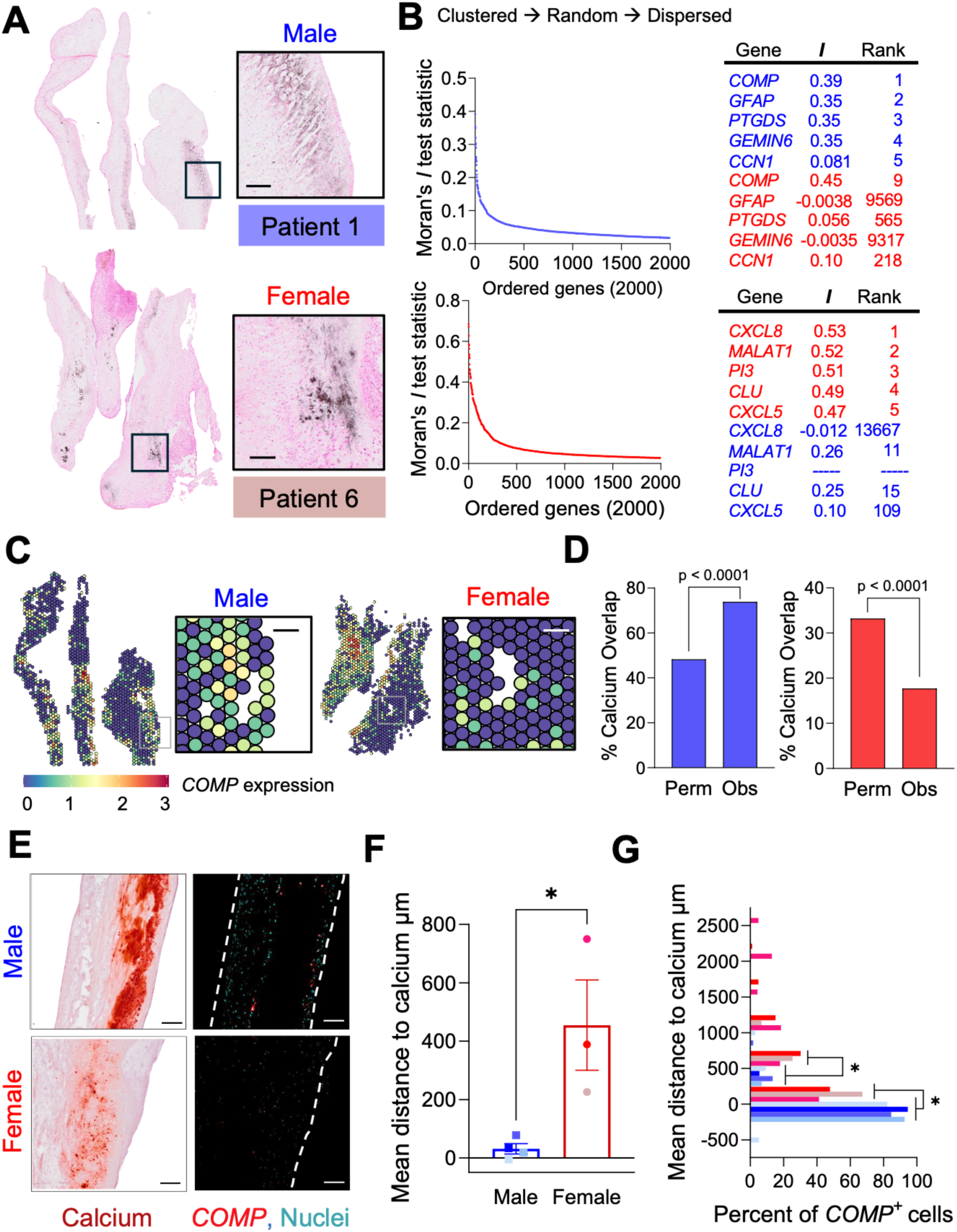
*COMP* gene expression overlaps with calcium in male tissue. **A.** Representative images of Von Kossa-stained tissues from patient 1 (male) and patient 6 (female). Scale bars = 100 µm. **B.** Moran’s *I* test statistic for the top 2000 genes and Moran’s *I* of male and female differentially expressed genes ranked by order. **C.** Spatial transcriptomics spot expression for *COMP* gene. Scale bars = 100 µm. **D.** Permutated and observed calcium overlap for male and female spatial transcriptomics maps. Significance determined by Fisher’s exact test. **E.** Representative images of Alizarin red stained tissues with corresponding RNA-FISH images of diseased male and female tissues of *COMP*^+^ cells near sites of calcification. Scale bars = 100 µm. **F.** Mean distance of *COMP*^+^ cells to sites of calcification for subset of male diseased (N = 4) and female diseased (N = 3) patients. Means ± standard deviations and *P-*values of comparisons shown with statistical significance determined via Welch’s t test. \**P* <0.05. **G.** Frequency distribution of percentage of *COMP*^+^ cells at binned distances from calcification; Means shown; \**P* <0.05.

We chose to further investigate and validate *COMP* as a potential regulator of male-biased valve calcification. Our rationale was three-fold: (1) *COMP* was our top ranked clustered gene in male tissue, (2) *COMP* had male-biased expression in the most abundant VIC 1 subcluster, and (3) previous literature reporting *COMP* is upregulated in diseased aortic valves and atherosclerotic plaques^1,37^. Our spatial transcriptomics analysis revealed high overall mapping of *COMP* expression spots in male and female tissues (**Figure 4C**). A Monte Carlo analysis of observed *COMP* localization to calcified regions versus randomly permutated overlap shows *COMP* overlaps with calcification specifically in male tissue relative to female tissue (\*\*\*\**P*<0.0001) (**Figure 4D**). We next utilized RNA-FISH staining to validate male-specific *COMP* localization near calcification sites. The mean shortest distance between *COMP* expressing nuclei and regions of calcification in tissue was significantly shorter in male tissues (31.3 ± 35.1 μm) relative to female tissues (455.0 ± 268.2 μm) (**Figure 4E – 4G**). The percentage of *COMP* positive nuclei trends towards increased expression in diseased male tissue (10.3 ± 4.4%) relative to diseased female tissue (4.0 ± 3.8%) (**Supplementary Figure VII**). Thus, we verified that *COMP* expression in male tissue localizes near sites of calcification across multiple patient samples.

### Macrophage sub clustering analysis reveals female specific increase in antigen presenting macrophages

We next determined female-biased, inflammation-driven modulation of valve calcification, with the hypothesis that macrophage-VIC interactions modulate increased fibrotic thickening and decreased calcium surface area in female valves. First, we investigated CD68+ macrophage presence across sex-separated healthy and diseased aortic valve tissue. Whole tissue immunofluorescence staining for CD68 revealed diseased aortic valve tissue had patient-dependent increases in CD68+ macrophages relative to healthy patients (**Figure 5A**), and female diseased tissues had a ∼1.9-fold increase in CD68+ macrophages relative to male (**Figure 5B**). We observed a positive correlation between percentage of CD68+ macrophages and percentage of calcium over surface area in males (R^2^ = 0.8992, *P* = 0.0024), relative to female tissue that showed no correlation between CD68+ macrophages and calcified area (R^2^ = -0.2082, *P* = 0.59) (**Supplementary Figure VIII)**.

**Figure 5.**
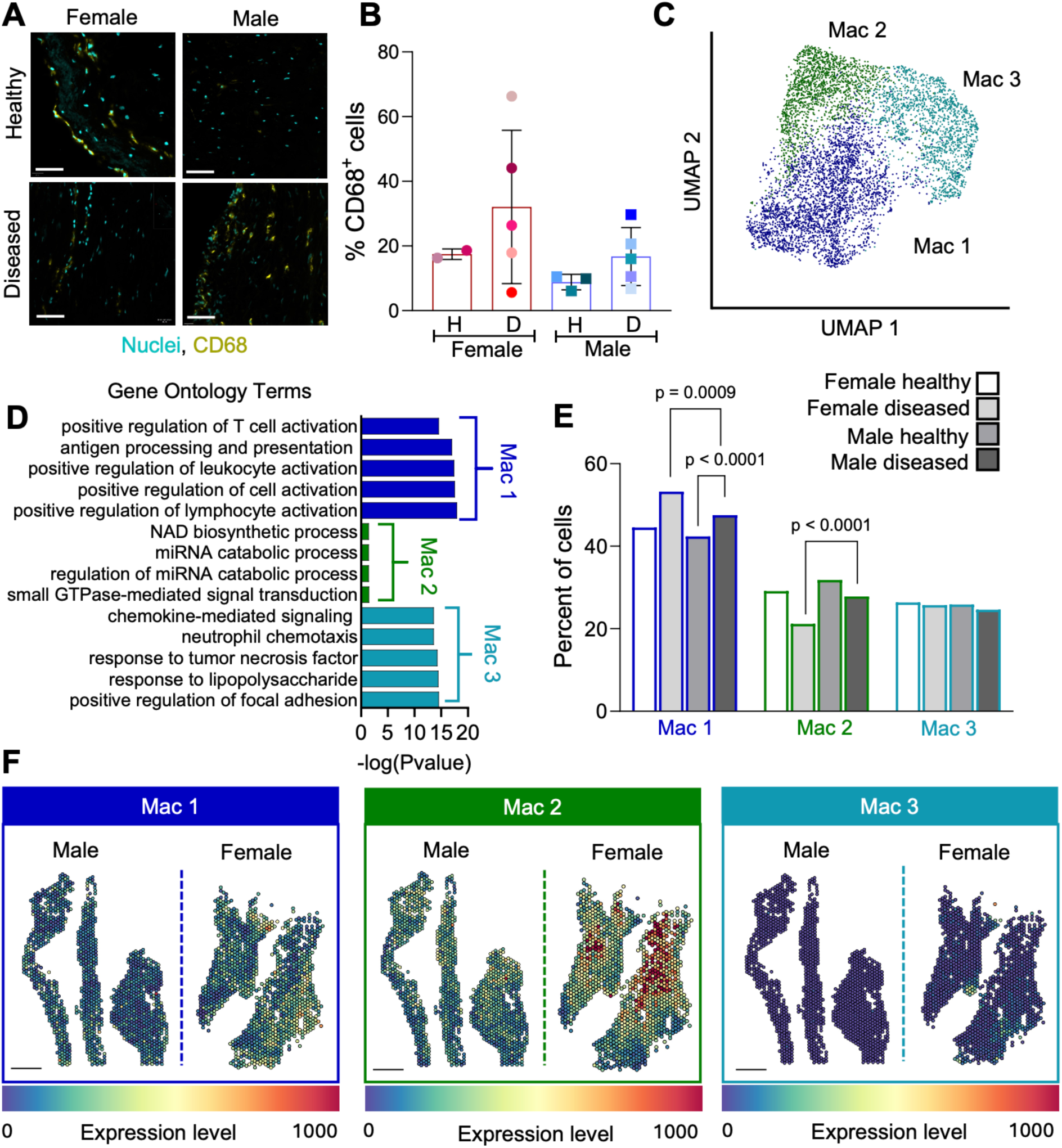
Macrophage sub clustering analysis reveals female-specific increase in antigen presenting macrophages. **A.** Representative images of CD68 staining colocalized with cell nuclei in male and female valve tissue. Scale bars = 100 µm. **B.** Percentage of CD68 positive cells in sex separated valve tissue; female healthy (N = 2), female diseased (N = 5), male healthy (N = 3), male diseased (N = 5); Means ± standard deviations and *P*-values of comparisons shown with statistical significance determined via two-way ANOVA with Tukey posttests. **C.** UMAP of subclustered VIC populations **D.** Top 5 Gene ontology terms for each macrophage subtype. **E.** Proportions of VIC subpopulations for female healthy (N = 1), female diseased (N = 3), male healthy (N = 4), male diseased (N = 4). *P-*values of comparisons shown with statistical significance determined via chi-squared analysis. **F.** Top 5 single cell RNA sequencing gene marker for each VIC subtype scored onto spatial transcriptomics maps. Scale bar = 1000 μm.

We chose to further investigate macrophage subtypes that may exhibit sex-specific gene expression signatures associated with fibro-calcification, given sex differences in correlations between macrophage presence and calcification. We observed 3 macrophage subclusters in our male and female datasets with the following upregulated expression markers: Mac 1 (*C1QB, C1QA, CTSB, FTL, C1QC*), Mac 2 (*MALAT1, DOCK4, AKAP13, FNIP2, NEAT1*), and Mac 3 (*CCL3, CCL4, CCL3L1, CXCL3, IL1B*) (**Supplementary Figure IXA**). Differential expression analysis between male and female diseased macrophage populations shows that *S100A10, HLA-DRB1, CXCL5* are female biased while *CCL20*, *CXCL1*, *CXCL2* are the top male biased genes (*P* < 0.0001, log_2_FoldChange > 0.5), with associated sex-biased pathways (**Supplementary Figure IXB – IXC)**. Additionally, we observed *SPP1* and *ACTB* are uniquely biased in the female Mac 3 population. Significant GO terms associated with Mac 1 include antigen presentation and T cell recruitment whereas Mac 3 expresses chemokines and chemokine ligands (*P* < 0.0001) (**Figure 5D**). Mac 2 is associated with GO terms microRNA catabolic process and NAD biosynthetic process (*P* < 0.0001). Consistent with T cell recruitment markers in Mac 1, we observe T cell marker positive spots with male and female tissue (**Figure 2H**).

We then determined the distribution of each macrophage subtype across sex and found the proportion of Mac 1 is higher in diseased females than in diseased males (χ^2^ = 11.06, 1 df, *P* = 0.0009). Additionally, we observed a higher proportion of Mac 1 in diseased females relative to healthy females (χ^2^ = 16.68, 1 df, *P* < 0.0001). Lastly, we observe a higher proportion of Mac 2 in diseased males relative to diseased females (χ^2^ = 21.72, 1 df, *P* < 0.0001). We next visualized the distribution of macrophage heterogeneity across the tissue by scoring the top 5 markers for each subtype. We observed expression of markers associated with all macrophage subtypes (**Figure 5F**). Within the female population, we observe spatial heterogeneity of macrophage subtypes and colocalization of Mac 1 and Mac 3 populations. Taken together, we revealed macrophage subtypes that may promote sex-dependent signaling networks associated with antigen presentation, microRNA degradation, and response to pro-inflammatory chemokines.

### Cell communication analysis reveals increased macrophage to VIC communication via SPP1-CD44 in female tissue

We next investigated sex differences in intercellular interactions that drive sex-dependent gene expression changes in aortic valve tissue. Using Cell Chat, we investigated ligand-receptor binding between macrophages, VICs, VECs, and T cells using our single cell sequencing dataset (**Supplementary Figure X)**. Focusing on macrophage interactions with VICs, VECs, and T cells, we observed significant interactions between osteopontin (*SPP1*, also known as Secreted Phosphoprotein 1 or osteopontin) from macrophages and other receptor signaling pathways such as CD44 and subunits alpha V and beta I of the integrin protein specifically among female VIC, T cell, and VEC populations (**Figure 6A**). On the other hand, we observe male-specific macrophage ligand – VIC receptor interactions, including PPIA to BSG, NAMPT to integrin subunits alpha V and beta I, and MIF to ACK3 (**Figure 6B)**.

**Figure 6.**
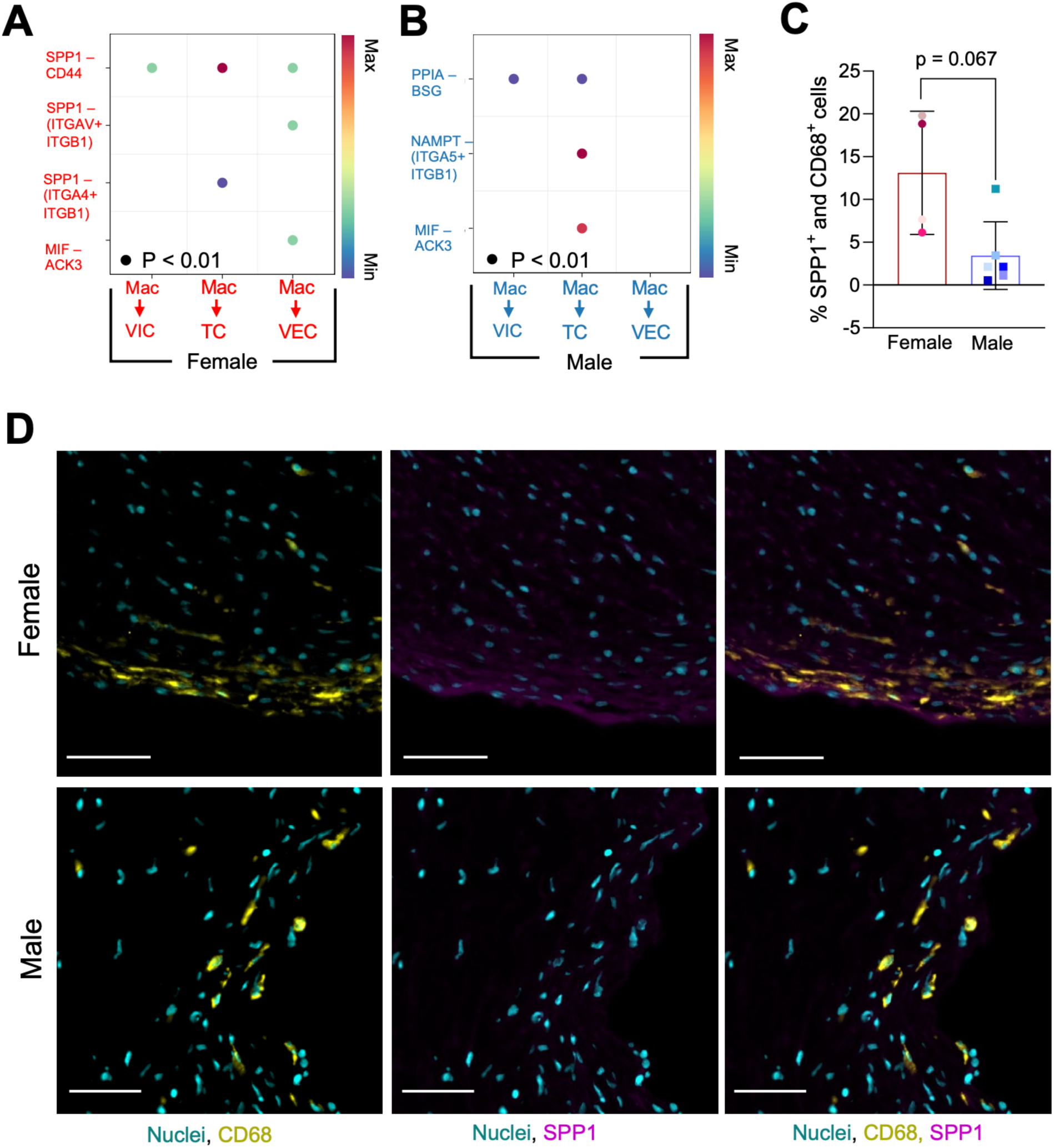
Female biased SPP1^+^ macrophage signaling and presence in valve tissue. **A.** Ligand – receptor cellular communication probability for single cell RNA seq; female healthy (N = 1), female diseased (N = 3). *P* < 0.01 indicates a significant interaction. **B.** Ligand – receptor cellular communication probability for single cell RNA seq; male healthy (N = 4), male diseased (N = 4). *P* < 0.01 indicates a significant interaction. **C.** Percentage of macrophage cell subtypes expressing SPP1 for female diseased (N = 4), male diseased (N = 6) via protein staining. Means ± standard deviations and *P-*values of comparisons shown with statistical significance determined via Welch’s t test. \**P* < 0.05. **D.** Representative images of SPP1 and CD68 protein staining in male and female valve tissue. Scale bar = 50 μm.

Next, we sought to quantify macrophage protein expression of SPP1 in male and female valve tissue. We observed an increased percentage of SPP1^+^ and CD68^+^ macrophages in female (13.11 ± 7.20%) relative to male (3.44 ± 3.95%) diseased tissue relative to male diseased tissue, which trended towards significance (**Figure 6C**). Representative images reveal colocalization of SPP1 and CD68 expression in female tissues relative to male tissues (**Figure 6D**). At the transcriptional level we quantified the percentage of *CD44*^+^ VIC subtype expression and found no significant differences between patient groups (**Supplementary Figure XI**). However, we observed an increase in *CD44* expression in VIC 4, a VIC subtype expressing biomarkers for immune cell interaction.

As a strategy to define “neighborhoods” of cells within the tissue, we utilized unsupervised clustering of spatial transcriptomics RNA spots (**Figure 7A**). We identified 6 cluster groups residing within the tissue: g1 (*PRELP, CLU, TIMP3, MGP, VCAN*), g2 (*CXCL5, CXCL8, HLA-B, FTH1, SPP1*), g3 (*COMP, PRELP, AEBP1, LTBP2, FMOD*), g4 (*C7, CCN1, PTGDS, C11orf96, THBS1*), g5 (*AL627171.2, CFH, ID4, DCN, NR4A1*), and g6 (*C7, CFH, GSN, MGP, TIMP3*) (**Supplementary Figure XII**). Focusing on spatial transcriptomics RNA clusters that overlap with macrophage subtypes, we observed a 17.9% overlap and 4% overlap of gene expression between g2 and Mac 1 and Mac 3, and no overlap with Mac 2 (**Figure 7B**).

**Figure 7.**
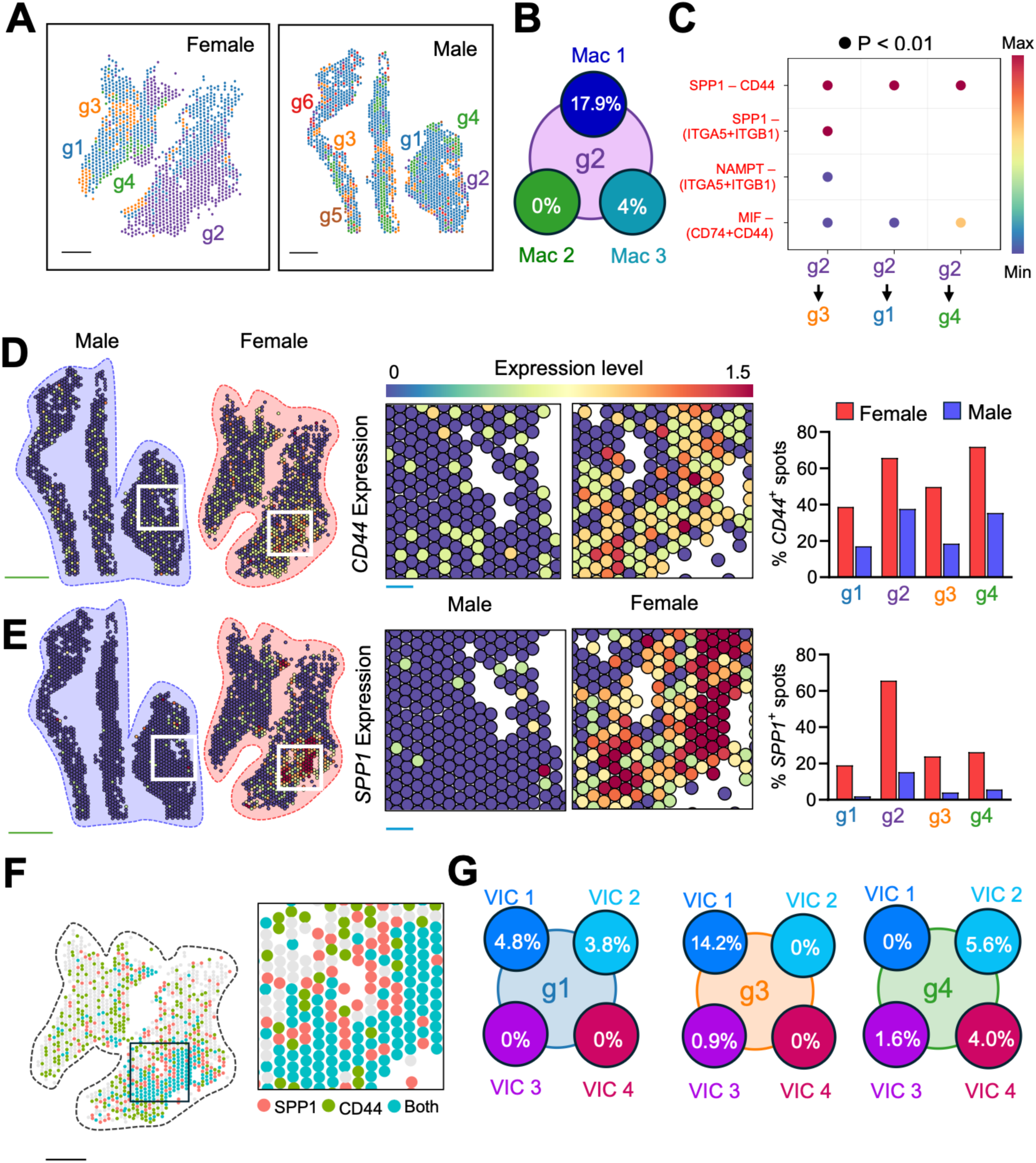
Female biased SPP1 – CD44 macrophage to fibroblast ligand binding in female tissue. **A.** Unsupervised clustering of spatial transcriptomics clusters. Scale bar = 1000 μm. **B.** Percent overlap of spatial transcriptomics RNA cluster g2 and single cell populations Mac 1, Mac 2, and Mac 3. **C.** Ligand – receptor cellular communication probability for spatial transcriptomics RNA clusters. *P* < 0.01 indicates a significant interaction**. D.** Spatial transcriptomics spot expression for *CD44* and percentage of positive spots for each spatial RNA cluster. Green scale bar = 1000 μm. Blue Scale bar = 100 μm. **E.** Spatial transcriptomics spot expression for *SPP1* and percentage of positive spots for each spatial RNA cluster. Green scale bar = 1000 μm. Blue Scale bar = 100 μm. **F.** Spatial transcriptomics spot expression for *SPP1* and *CD44* genes in female tissue. Scale bar = 1000 μm. **G.** Percentage overlap between single cell populations VIC 1, VIC 2, and VIC 3 and spatial transcriptomics RNA clusters g1, g3, g4.

We next validated our Cell Chat analysis with our spatial transcriptomics dataset and hypothesized that female valve tissue would contain overlapping *CD44* and *SPP1* expression spots. We observed g2 exhibited female tissue specific communication probability of *SPP1* with *CD44* and integrin subunits alpha V and beta I (**Figure 7C**). We observed no significant communication probability between g2 and other spatial transcriptomics clusters in male tissue. Furthermore, we observed an increased expression percentage of *SPP1* and *CD44* positive spots in female tissue relative to male tissue (**Figure 7D – 7E**). We also observed high co-localization of *SPP1* and *CD44* in female tissue, whereas male tissue had no co-localization of *SPP1* and *CD44* (**Figure 7F**). We also observed *SPP1*^+^ macrophages act on spatial transcriptomics clusters g1, g3, and g4 and sought to identify overlap between targeted clusters and VIC subtypes. We observed the highest overlap between VIC 1 and g3 (14.2%) and g1 (4.8%) while VIC 2 had the highest overlap with g4 (5.6%) (**Figure 7G**). As such, our analyses revealed sex-dependent cellular communication networks between VIC and macrophage populations that may modulate valve fibro-calcification.

## Discussion

Our data reveal sex-dependent gene expression signatures in VICs near sites of calcification using spatial transcriptomics analyses of human aortic valves. VICs have long been recognized as the primary modulators of aortic valve fibro-calcification as the most abundant cell population in the tissue. Indeed, VICs have been shown to modulate sex-dependent myofibroblast signaling pathways via X-chromosome^8^ and Y-chromosome^9^ linked genes, in addition to sex hormone associated signaling networks^19,38,39^. Our histological analyses of male and female tissues also corroborate numerous clinical studies revealing increased calcification burden in male AVS patients and reduced calcification in the early stages of AVS for female patients^4,6,18^. Our work adds to this increasing body of literature to reveal location-dependent and sex-specific gene expression signatures in heterogeneous populations of VICs that orchestrate aortic valve calcification. We posit the VIC populations near sites of calcification should be the first targets for pharmacological intervention to prevent or halt further AVS progression.

We have identified and validated COMP expression in male VICs near sites of calcification, implicating *COMP* as a male-specific regulator of mineralization and subsequent valve calcification. Collagen oligomeric matrix protein (*COMP*), is a key regulator of chondrogenesis and bone growth and has also been linked to progression in various cardiovascular diseases^1,37,40^. In one example, *COMP* expression was increased in vascular smooth muscle cells that localized to plaques in symptomatic atherosclerosis patients^37^. Additionally, myofibroblast lineage tracing in a myocardial infarct mouse model revealed a *COMP* positive matrifibrocyte that persists in scar tissue^41^. Of note, these studies were carried out with only male samples, leaving a gap in knowledge regarding sex-dependencies in *COMP* expression and cardiovascular disease progression. To the best of our knowledge, our work is the first to demonstrate sex-dependent *COMP* expression and localization near sites of calcification in male aortic valves relative to female valves. We suspect localized *COMP* expression is epigenetically regulated in male VICs, based on previous studies revealing histone deacetylase 3 (HDAC3) inhibition promotes *COMP* expression in mesenchymal stem cells differentiating to chondrocytes. Another study demonstrates HDAC3 activity suppresses Runt-related transcription factor 2 translocation in preosteoblasts halting their differentiation to mature osteoblasts. Furthermore, Ubiquitously transcribed tetratricopeptide repeat, Y chromosome (UTY) is known to modulate male-specific osteoblast-like differentiation in VICs when exposed to nanoparticles that mimic sites of calcification^9^. UTY acts on histone H3 upstream of *COMP*^42^, suggesting that increased nano-scale matrix stiffness due to calcium particles may induce male-specific epigenetic modifications driving *COMP* gene expression. Future *in vitro* studies using valve matrix-mimicking hydrogel biomaterials^43^ will be beneficial to further investigate the role of male-biased epigenetic modifications and expression of *COMP* and other osteoblast-associated genes.

We also posit that AP1 in VICs is a critical modulator of male-dependent aortic valve calcification. VICs in our dataset exhibit a strong correlation with the FOS/JUN (AP-1) transcriptional activity across all male VIC populations relative to female VICs. AP-1 signaling is linked to osteoblast-like differentiation of VICs via Tumor Necrosis Factor (TNF) binding to the RANK receptor^36^. Additionally, p38 MAPK signaling regulates the transcription of FOS and JUN, thereby increasing the expression and activation of AP-1 complexes^44^. Once activated, AP-1 participates in transcriptional regulation of downstream genes linked to apoptosis, ECM remodeling, cellular differentiation, and immune cell infiltration, which are all hallmarks of AVS progression^45^. To the best of our knowledge, no prior studies have demonstrated sex differences in AP-1 signaling during AVS progression. Previous studies have revealed male-dependent p38 MAPK activity in myocardial ischemia^46^ and AVS progression^10^, further pointing to p38 MAPK regulation of AP-1 as a key modulator of male-dependent valve calcification, and may serve as potential male-biased target to mitigate VIC transdifferentiation into osteoblast-like cells that exacerbate calcification.

Our observations also lead us to suspect inflammatory macrophages are a major contributor to female-specific aortic valve thickening and subsequent fibrosis. Valve thickening is a hallmark of calcific aortic valve stenosis progression due to increased extracellular matrix deposition, calcium particle formation, and immune cell infiltration^21,31^ Although we did not observe increased collagen deposition in female valves as observed in other studies with greater patient cohort sizes^4^, we found increased valve thickening in our female valve leaflet samples, despite lower calcification area measurements. We have identified an ECM remodeling VIC population (VIC 1) which upregulates ECM protein coding genes including fibronectin and decorin in female valves^47^. Therefore, we suspect VIC 1 contributes to aberrant ECM deposition and aortic valve thickening. Furthermore, females have an increased proportion of antigen presenting macrophages (Mac 1), which secrete a myriad of cytokines that recruit other inflammatory cells, which may also contribute to valve thickening^21,48^. Indeed, prior work has also established that secreted factors from inflammatory macrophages contribute to VIC proliferation *in vitro*^49,50^. Our observations suggest that collagen fiber density alone may not be a consistent measure for valve fibrosis and may need to be coupled with additional measures including valve thickening, macrophage accumulation, glycosaminoglycan deposition, and expression of myofibroblast markers including periostin and tenascin-c^51^.

Our spatial transcriptomics analysis has also revealed female-dependent VIC-macrophage interactions via SPP1-CD44 that may reduce calcification burden in females. In the early stages of aortic valve calcification, osteopontin^52^ is highly expressed and can sequester calcium phosphate in early AVS stages to reduce calcification burden^53–55^. Female-dependent osteopontin expression has also been previously reported in human aortic valves^12^, pointing to osteopontin as a female-dependent mediator of reduced calcification burden. Using spatial transcriptomics, our work has revealed a pro-inflammatory macrophage subtype (Mac 1) that expresses *SPP1* in female valves, which may bind to *CD44*^56^ in female VICs to inhibit pro-calcification programs. Previous studies have shown that macrophage and fibroblast interaction can drive myofibroblast activation and downstream fibrotic remodeling^22,23^. Specifically, *SPP1*^+^ macrophage and *CD44* fibroblast interactions drive fibrosis across different organs, such as the lungs, skin, heart, and kidney^57–59^. We posit the crosstalk could lead to increased fibrotic burden in female valve tissue. We also suspect that SPP1-CD44 macrophage-VIC interactions may be synergistically driven via sex chromosomes and sex hormones. A previous study treated bovine coronary smooth muscle cells with β-estradiol in the presence of osteogenic medium resulted in decreased OPN^20^. Although the inhibitory effects of estrogen on valvular calcification are varied in literature, likely due to the synergistic contribution of sex chromosomes^8^ in disease progression, our observations partially corroborate observations of decreased calcification during disease progression. Further *in vitro* work utilizing estrogen and SPP1 dosing on VIC populations is required to further elucidate the role of SPP1 in calcium attenuation.

In conclusion, our work provides a blueprint for dissecting the sex-dependent cellular drivers of aortic valve fibro-calcification, in anticipation of future development of sex-specific treatments for AVS patients. Our study defines cellular heterogeneity in a sex-specific context, in which we observed spatial proximity of *COMP* to sites of calcification in males, contrasted with *SPP1-CD44* mediated macrophage-VIC signaling in females. Our findings suggest that sex differences in AVS progression are not merely systemic in scale but are also embedded in the local cellular microenvironment of the valve. Importantly, future efforts should prioritize targeting VIC populations near calcified regions for paths to treatment using combinations of small molecule inhibitors^60^. Our results underscore the need for future studies to determine sex-specific epigenetic and paracrine regulatory networks using additional spatial transcriptomics technologies. Together, our multi-omics approach underscores the importance of understanding valve cell heterogeneity in a spatial context when elucidating sex differences in valve disease and improving future patient outcomes with sex-based precision medicine.

## Methods

### Human aortic valve specimens

Healthy and diseased valve tissue specimens were obtained from cadaver hearts (N = 21 total patients) with postmortem intervals ranging from 36 to 48 hours through the National Disease Research Interchange as approved by the University of California San Diego (IRB 804209). Patient inclusion criteria were defined as significant risk factors for fibro-calcification such as Aortic Stenosis, Calcific Aortic Valve Disease, Diabetes Mellitus II, and Chronic Hypertension for more than 10 years (**Supplementary Table I**). We excluded patients with bicuspid aortic valve morphology, rheumatic aortic stenosis, and ineffective endocarditis. Human hearts were shipped on ice, aortic valves were excised, and valves were flash frozen in optical cutting temperature (OCT) for spatial transcriptomics or digested for single cell RNA sequencing. Sample sizes for all experimental procedures are provided in **Supplementary Table II**.

### Valve tissue histology

Patients were stratified into healthy and diseased cohorts based on quantified measures of calcified area in tissue and Warren-Yong scoring^31^. Frozen tissue blocks were sectioned at 10 µm and mounted on slides for subsequent staining. Alizarin Red (Abcam, Cat. No. 146374) and Von Kossa staining kits (Fisher Scientific, Cat. No. NC9239431) were used to visualize calcification. For Alizarin red staining, sections were equilibrated to room temperature for 10 minutes and incubated in 2% Alizarin Red solution adjusted to a pH of 4.1. Tissue sections were dipped 20 times in acetone, acetone-xylene (1:1), and xylene. Von Kossa staining was carried out as previously described^12^. To visualize collagen abundance, tissue sections were stained with Masson’s Trichrome (Sigma-Aldrich Cat. No. HT15-1KT). Briefly, sections were equilibrated to room temperature for 10 minutes and fixed in an acetone-methanol solution (60:40) at -20°C for 10 minutes and rinsed under running tap water for 5 minutes. Sections were then immersed in Bouin’s solution (Sigma-Aldrich, HT101128) for 15 minutes at 56°C, cooled in tap water at room temperature (18°C-26°C), and subsequently washed in running tap water for 5 minutes. Next, slides were immersed in a Weigart Iron hematoxylin solution for 5 minutes, washed with running tap water for 5 minutes and rinsed in deionized water. Sections were incubated in a Biebrich Scarlet-Acid Fuchsin for 5 minutes and briefly rinsed in deionized water. Slides were then placed in phosphotungstic-phosphomolybdic acid solution (1:1) for 5 minutes and directly immersed in an aniline blue solution for 5 minutes. Tissues were immediately placed in 1% acetic acid solution for 2 minutes. Finally, slides were rinsed in tap water, dehydrated through increasing concentrations of ethanol and cleared in xylene. All tissues were mounted in VectaMount mounting medium (Fisher Scientific, Cat. No. NC2004138) for subsequent imaging. Whole tissue imaging was carried out on the Olympus VS200 Slide Scanner with a 20X objective.

### Histological analysis

Images of tissue sections were analyzed in Qupath^30^. To quantify intact calcium surface area, a Random Tree Algorithm was used to train a pixel classifier to detect regions of calcium which were normalized to tissue area. Only valve sections with intact calcium deposits were included for quantification. To quantify tissue fibrotic area, a pixel classifier was trained on Masson’s trichrome stained tissue. Dark blue pixels were defined as dense fibers and normalized to whole tissue area. We utilized previously reported Warren-Yong scoring to classify patients as healthy and diseased^31^.

### Scanning Electron Microscopy (SEM) and calcium particle size quantification

Fresh frozen human valve leaflets were cut in 10 µm sections onto glass slides before fixation with 4% paraformaldehyde for 10 minutes at room temperature. Specimens were serially dehydrated by washing for 1 hour in increasing concentrations of ethanol diluted in DPBS (10% to 100%). After the final dehydration, specimens were set to dry overnight before iridium sputter coating for 40 seconds at 75% power. Specimens were imaged using a scanning electron microscope (FEI Quanta FEG 250) using manufacturer settings. For each patient, at least three calcific regions of interest were imaged for quantification as determined by Alizarin Red staining. Calcium particle size was quantified via ImageJ as previously described^9^.

### Isolation of human valve cells for single cell sequencing

Aortic valve leaflets were excised from the aortic valve by cutting along the valve base and rinsed with Earle’s Balanced Salt Solution (EBSS) (Gibco, Cat. No. 24010043). All 3 leaflets per specimen were cut in half with 3 halves placed on ice for spatial transcriptomics protocols and three halves digested for single cell isolation and sequencing. Tissues were minced in Liberase TM (Sigma-Aldrich, Cat. No. 5401119001) in EBSS at 0.25 WU/mL and digested on an orbital shaker at 175 RPM for 30 mins at 37°C. DNAse I (Sigma Aldrich, Cat. No. DN25) was added to digestion medium at 100 µg/µL to prevent cell aggregation from genomic DNA. After digestion cells were immediately quenched with 10% Fetal Bovine Serum (FBS, Thermo Scientific, Cat. No. 16000069) in Medium 199 (Thermo Fisher Scientific, Cat. No. 11043023). Cells were strained in 3 successive straining steps with 100 µm, 70 µm, and 30 µm strainers and centrifuged at 400 rcf for 10 minutes. The cell pellet was resuspended in red blood cell lysis buffer (Thermo Scientific, Cat. No. 00433357) and rinsed using 10% FBS in media. After a final centrifugation step, cells were resuspended in 200 µL of 10% FBS media and counted using a cell hemacytometer prior to cell encapsulation in gel-beads for single cell RNA sequencing.

### Single cell sequencing

Aortic valve leaflets (N = 7 patients from our cohort) were analyzed using single cell sequencing. 2000-5000 cells per sample were targeted on the Chromium platform (10X Genomics) using a single lane per patient. Single cell mRNA libraries were built using the Chromium Next GEM Single Cell 3’ Library Construction V3 Kit and libraries were sequenced on the Illumina NovaSeq 6000. Sequencing results statistics and quality control are summarized in **Supplementary Tables III – V**. The *Homo Sapiens* genome and gene annotations (assembly: GRCh38) were used to align data via the Cell Ranger 8.0.0 (10X Genomics) “count” command to generate expression matrices. Our data were integrated with previously published data sets^26^ (N = 5 patients) for a total of 12 patients for subsequent data processing.

### Single cell RNA sequencing processing

Downstream analysis on patient gene expression matrices was performed using the Seurat^32^ v4.4.0 pipeline in R. Matrices were filtered from dead cells and cells with high (> 7000) and low (< 200) feature counts. Normalization was performed using Seurat’s SCT transform v.2 which applies a negative binomial regression model to correct for technical artifacts such as sequencing depth variation while maintaining highly variable biological features. The normalized data was then integrated using Pearson’s residuals via the functions PrepSCTIntegration() followed by FindIntegrationAnchors() and IntegrateData() in Seurat. Dimensional reduction was performed on the integrated data set with the RunPCA() and RunUMAP() functions with 30 principal components. Graph-based clustering was performed using *k-nearest* neighbor FindNeighbors() and transformed into clusters using FindClusters() at a resolution of 0.01 on the data set. Wilcoxon rank sum testing was used to identify differentially expressed genes between different patient groups. Gene Ontology enrichment of differentially expressed genes was performed using enrichGO R-4.4.2, upstream regulator enrichment was performed using Ingenuity Pathway Analysis (Qiagen), and pathway analysis was performed using EnrichR^33^.

### Spatial transcriptomics

Aortic valve leaflets embedded in OCT medium were cryosectioned at 10 μm in thickness at -20°C. Spatial transcriptomics protocols (10X Genomics) were performed on tissue sections with an RNA Integrity Number (RIN) > 7 measured on the Agilent TapeStation for all samples. Sections were stained with Hematoxylin and Eosin as described in 10X Genomics protocol and imaged at 20X magnification on the Nikon Ti2-E widefield microscope. The tissue sections were then processed for spatially resolved gene expression according to protocols from the 10X genomics Visium Spatial Transcriptomics kit. The tissue was permeabilized for a previously optimized time of 24 minutes to release mRNA for reverse transcription into cDNA. Sequencing libraries were prepared and assessed for quality control on the Agilent Tapestation prior to sequencing on the NovaSeq 6000 instrument. The resulting sequencing data was aligned to images using SpaceRanger v.3.0.0 (10X Genomics) to generate gene expression matrices for subsequent analysis and processing. Sequencing results statistics and quality control are summarized in (**Supplementary Tables VI – VIII).**

### Spatial transcriptomics processing

Spatial transcriptomics analysis was performed using the Seurat v4.4.0 pipeline in R. Gene expression matrices were initially filtered from high or low feature counts and regions of dead cells. Regions of calcification were confirmed via Von Kossa staining on tissue serial sections. Normalization was performed using Seurat’s SCT transform v.2 and merged our male and female replicates (N = 2) data into a single Seurat object and applied dimensional reduction and principal component analysis on the merged group. Then, male and female groups were integrated for comparison using Pearson residuals, FindIntegrationAnchors(), followed by PrepSCTIntegration(), and IntegrateData(). Graph-based clustering using *k-nearest* neighbor function FindNeighbors() and transformed into clusters using FindClusters() at a resolution of 0.1 on the entire integrated dataset. Wilcoxon rank-sum testing was used to identify differentially expressed genes between male and female samples.

### Moran’s I test statistic and calcium colocalization

The spatial distribution of genes was examined using the Moran’s *I* test statistic, which is a spatial autocorrelation coefficient used to quantify similar distribution of spatially clustered gene expression. Moran’s *I* ranges from 0 (homogenously distributed throughout sample) to 1 (significant spatial enrichment). Moran’s *I* was computed for each Visium spot (55 µm) which reveals genes that are spatially enriched and are likely expressed by cells in the same vicinity. To determine the probability of overlap between genes of interest and calcification, we ran a Monte Carlo simulation (n = 10,000) to randomly sample spatial transcriptomics spots across tissue samples and quantified the percentage of *COMP* positive spots overlapping with calcium versus random distribution.

### RNA Fluorescence In Situ Hybridization

RNA in situ hybridization was carried out on aortic valve leaflet sections (N = 7 patients) according to manufacturer protocols using the RNAscope Multiplex Fluorescent Reagent v2 (Advanced Cell Diagnostics Bio, Cat. No. 323100) kit. Probes for *COMP* (ACDBio, Cat. No. 457081) and *UBC* (Ubiquitin C, positive control to confirm RNA integrity, ACDBio, Cat. No. 320861) were used. As a negative control we used a probe that targets the *dapB* gene (ACDBio, Cat. No. 320871). Tissue sections at 10 µm were fixed in 4% PFA as listed above for 90 minutes and dehydrated in 50%, 70% and 100% ethanol. Samples were then blocked with hydrogen peroxide (ACDBio, Cat. No. 322381) for 10 minutes at room and treated with Protease IV (ACDBio, Cat. No. 322340) for 20 minutes at room temperature. Tissues were then hybridized with probes for *UBC*, *dapB* and *COMP* for 120 minutes and visualized fluorescently with TSA Vivid fluorophores (Tocris, Cat. No. 320871) and DAPI (Sigma-Alrich, Cat. No. 10236276001). Whole tissue scans were taken under a 20X dry objective on the Olympus VS200 Slide Scanner. *COMP* fluorescent signal was adjusted with positive *UBC* and a negative control *dapB* for each sample.

### RNA-FISH analysis

The percentage of *COMP* positive cells and distance of *COMP* positive cells to sites of calcification was quantified in Qupath. Briefly, tissue nuclei were detected with DAPI stain using the Cell Detection function. Nuclei overlapping with *COMP* fluorescence greater than 600 units of fluorescence were defined as positive cells using the Positive Cell Detection function. Regions of calcification were determined via serial sections stained with Alizarin Red and annotated on the tissue. The signed distance function was used to measure the shortest distance between *COMP* positive cells and regions of calcification in male and female tissues.

### Immunofluorescence staining

Aortic valve leaflets were analyzed using immunofluorescence staining techniques (N = 15). Briefly, tissue sections were brought to room temperature and then fixed for 12 minutes in 4% paraformaldehyde (Fisher Scientific, Cat. No. 50980487) solution in phosphate buffered saline (PBS). Tissues were outlined with a PAP pen (Abcam, Cat. No. 2601), permeabilized with 0.1% TritonX-100 (Fisher Scientific, ACS215682500) in PBS for 30 mins at room temperature and blocked in 5% bovine serum albumin (Sigma-Aldrich, Cat. No. A8327) and 4% Normal Goat Serum (Jackson ImmunoResearch, Cat. No. 005000121) in PBS for 1 hour at room temperature. Primary antibodies for osteopontin (Abcam, Cat. No. ab8448), CD68 (Abcam, Cat. No. ab955) and CD44 (Biotechne, Cat. No. AF6127) were diluted in blocking buffer and incubated overnight in a humidity chamber at 4°C. Samples were then equilibrated to room temperature and rinsed once 5 minutes in 0.1% Tween20 (Sigma-Aldrich, Cat. No. P1379) in PBS and then 2 times for 5 minutes in PBS. Samples were incubated in secondary antibodies AF647 (Invitrogen, Cat. No. A21247), AF555 (Invitrogen, Cat. No. A32727) and AF488 (Invitrogen, Cat. No. A11015) diluted in PBS for 1 hour in a humidity chamber at room temperature. Samples were rinsed 3 x 5 mins in PBS and mounted in ProLong Gold Antifade Mountant (Fisher Scientific, Cat. No. P36931) prior to imaging.

### Cell communication analysis

Cell Chat^34^, an R package, was used to determine ligand and receptor binding between single cell RNA sequencing populations using previously described methods^35^. Briefly, we utilized the IdentifyOverExpressionInteractions() function on the preprocessed single cell RNA sequencing object. Next, we computed the communication probability using the computeCommunProb() function using the trimean method. The communication probabilities of all ligand-receptor interactions were summarized into signaling pathways using the computeCommunProbPathway(). Cell to cell communication pathways with communication probabilities less than *P-*value of 0.01 were visualized with bubble plots.

### Statistical analysis

Statistical analyses were computed in GraphPad Prism (10.5.0) unless otherwise noted. Correlations were computed using two-tailed Pearson’s correlation with a 95% confidence interval and a *P-*value less than 0.05 was considered significant. Multiple comparisons testing was done to determine sex differences between healthy and diseased groups using a two-way ANOVA with Tukey posttests. Welch’s t tests were used to compute sex differences between male and female diseased cohorts and a *P-*value less 0.05 determined significance. Fisher’s exact testing was used to compute the difference between permutated and observed gene overlap with regions of calcification as well as sex differences in risk factors within the patient cohort. Significance of cell proportions for single cell RNA sequencing data were determined via Chi Squared analysis and only *P-*values less than 0.0009 were considered significant. Outliers greater than 3 standard deviations from the mean were removed to minimize influence of extreme values due to technical errors.

## Data availability

All data can be found in the manuscript and supplementary data sections of this manuscript.

## Acknowledgements

The authors would like to thank the patients that donated their organs and tissues to research. The authors acknowledge technical, sequencing, and research support provided by Dr. Trevor Biddle at the gCore in the Sanford Consortium for Regenerative Medicine and Dr. Kristen Jepsen at the Institute for Genomic Medicine in University of California San Diego. Additionally, the authors acknowledge the technical guidance provided by Dr. Elsa Molina and Cristian Quintero at the Single-Cell and Spatial Omics core in the Salk Institute for Biological studies. The authors would also like to acknowledge Zhenxing Fu who assisted with the preparation of human aortic valve tissue for spatial transcriptomics. The authors also acknowledge Zbigniew Mikulski at the La Jolla Institute for Immunology for guidance on tissue staining, imaging, and analysis. Lastly, the authors acknowledge Dr. Peng Guo and Dr. Richard Sanchez for their guidance on imaging of RNA-FISH tissues at the UCSD Nikon Imaging Center.

## Author Contributions

T.B. and B.A.A conceived and supervised the study. T.B and R.P. performed tissue histology and image analysis. R.M.G. performed all scanning electron microscopy and analysis. T.B., V.K.N., and R.P. and carried out single cell and spatial transcriptomics workflows. T.B. and V.K.N analyzed all transcriptomics data and subsequent statistical analyses. B.A.A and K.R.K. supervised all transcriptomics workflows and analyses. T.B. and B.A.A wrote and edited the manuscript. All authors approved the manuscript.

## Sources of funding

National Institutes of Health Pathway to Independence Award R00 HL148542 (BAA)

National Institutes of Health Director’s New Innovator Award DP2 HL173948 (BAA)

Chan Zuckerberg Initiative Science Diversity Leadership Award DAF2022-309430 (BAA)

American Heart Association 942253 (BAA)

National Science Foundation CAREER 2442606 (BAA)

National Institute of Health DP2 AR075321 (KRK)

National Institute of Health NHLBI T32HL007444 (VKN)

## Materials and correspondence

All correspondence should be addressed to Brian Aguado.

## Competing interests

No competing interests.

## Notes

https://www.ncbi.nlm.nih.gov/bioproject/PRJNA562645/

### Competing Interest Statement

The authors have declared no competing interest.

