## Supplementary Materials for "Spatial transcriptomic profiling of the human aortic valve reveals cellular sex differences near sites of calcification"

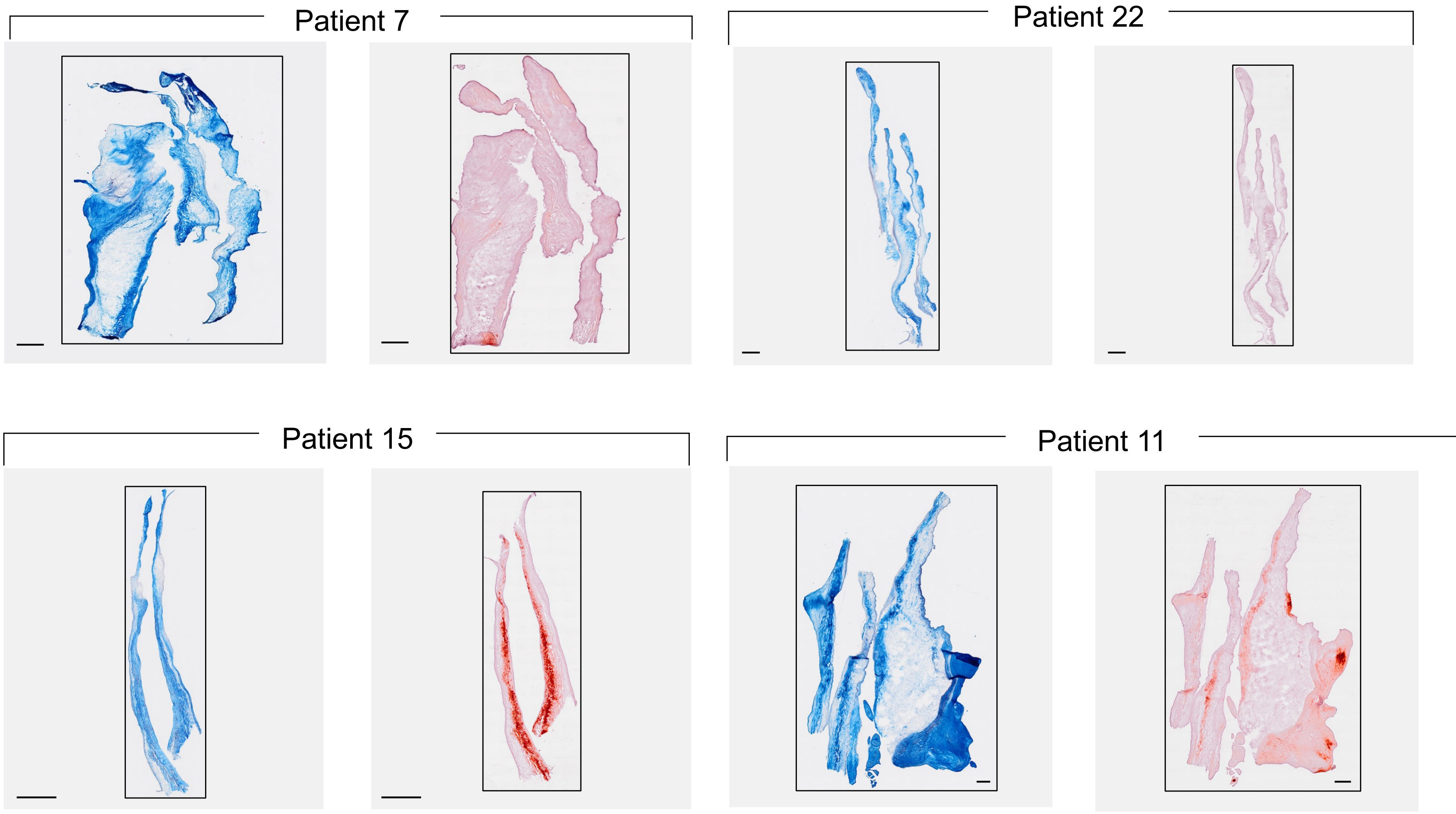


**Fig. I: Representative images of Masson's trichrome and Alizarin red staining of aortic valve leaflet sections.** Scale Bar = 500 µm.

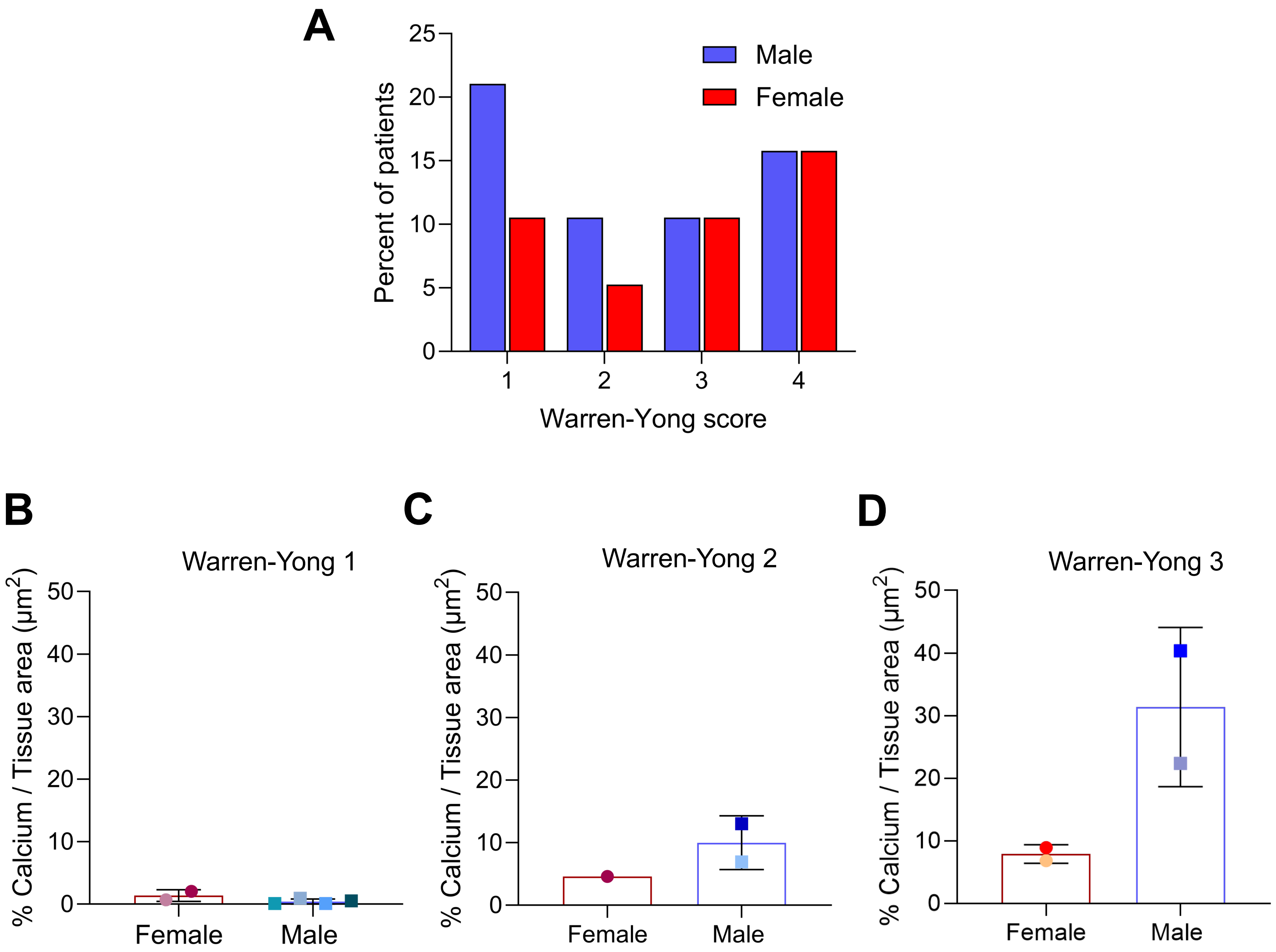


**Fig. II: Semi-quantitative Warren-Yong scoring of male and female tissue.** (A) Percent of male and female aortic valve tissues per Warren-Yong disease category. (B) Percentage of calcification over tissue surface area for a Warren-Yong score of 1. Sample size: N = 2 female tissues and N = 4 male with at least 2 leaflets per patient. Mean ± S.D. shown. (C) Percentage of calcification over tissue surface area for a Warren-Yong score of 2. Sample size: N = 1 female tissues and N = 2 male with at least 2 leaflets per patient. Mean ± S.D. shown. (D) Sample size: N = 2 female tissues and N = 2 male with at least 2 leaflets per patient. Mean ± S.D. shown.

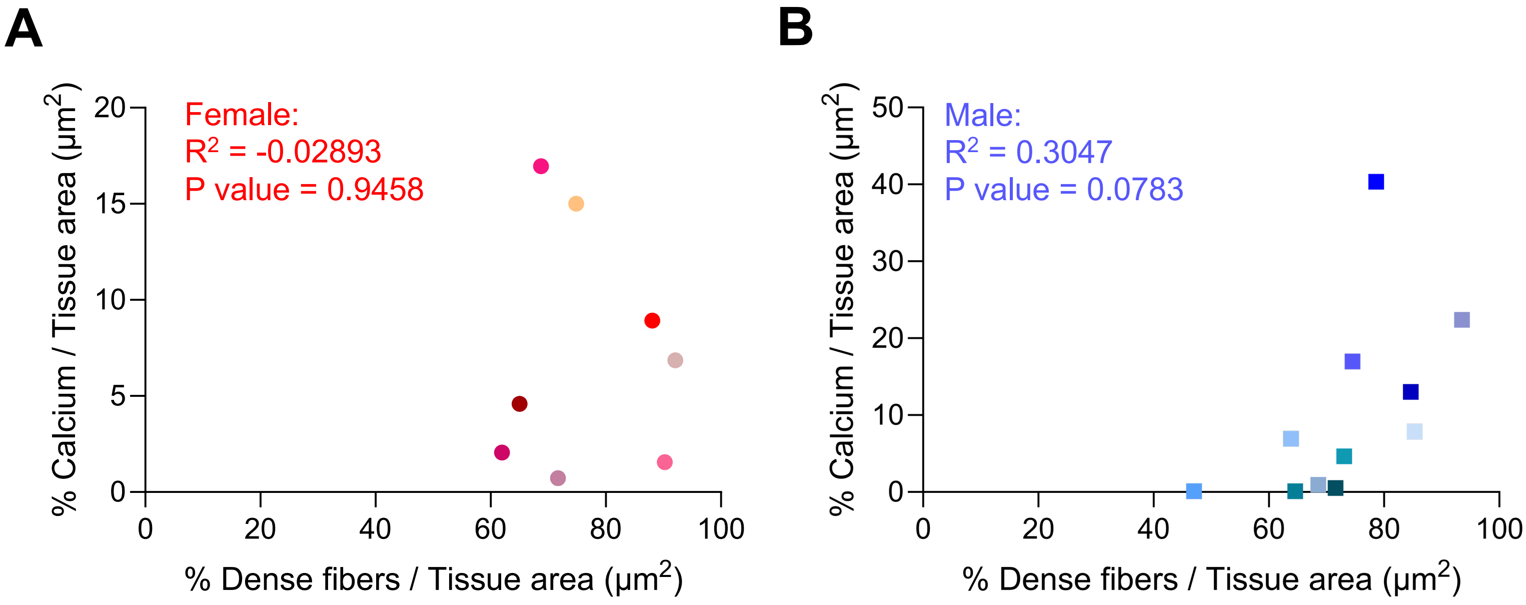


**Fig. III: Correlation of tissue calcification and fiber density in male and female valve tissue.** (A,B) Calcium over tissue surface area for female and male valve tissue specimens. Sample size: N = 8 female tissues and N = 11 male tissues with at least 2 leaflet per patient. Correlation tested with Pearson's correlation coefficient.

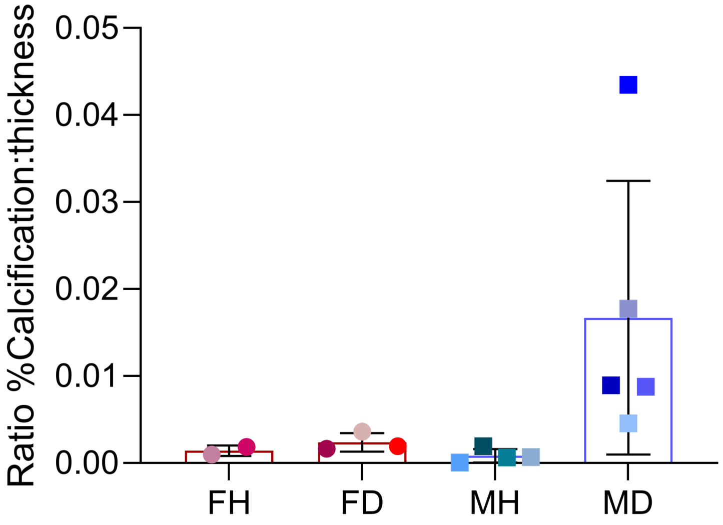


**Fig. IV: Ratio of percent calcification to tissue thickness.** Sample size: N = 2 female healthy tissues, N = 3 female diseased tissues, N = 4 male healthy tissues, N = 5 male diseased tissues with at least 2 leaflets per patient. Mean ± S.D. shown.

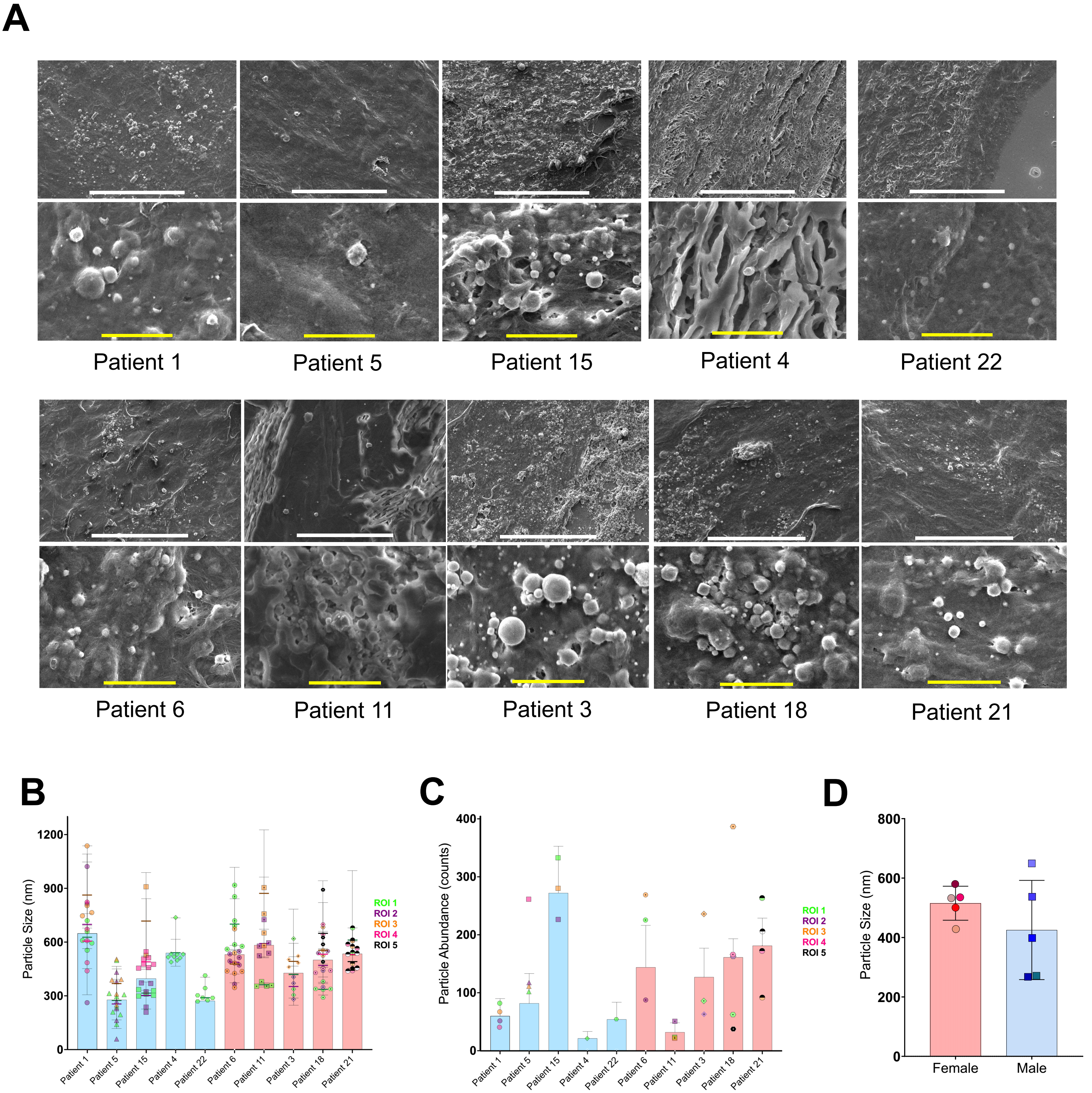


**Fig. V: Scanning electron microscopy of calcium particles in male and female tissue.** (A) Representative SEM images of human aortic valve tissue in male and female samples. White scale bar = 50 µm**.** Yellow scale bar = 5 µm. (B) Average particle size per region of interest in male and female human aortic valve tissue. Sample size: N = 5 biological replicates from 3 leaflet section with at least 3 images per region of interest. (C) Particle abundance in each ROI for male and female human aortic valve tissue. (D) Average calcium particle size in male and female tissue. Sample size: N = 5 female diseased tissues, N = 4 male diseased tissues. Significance tested with Welch's t-test (**P* < 0.05). Mean ± S.D. shown.

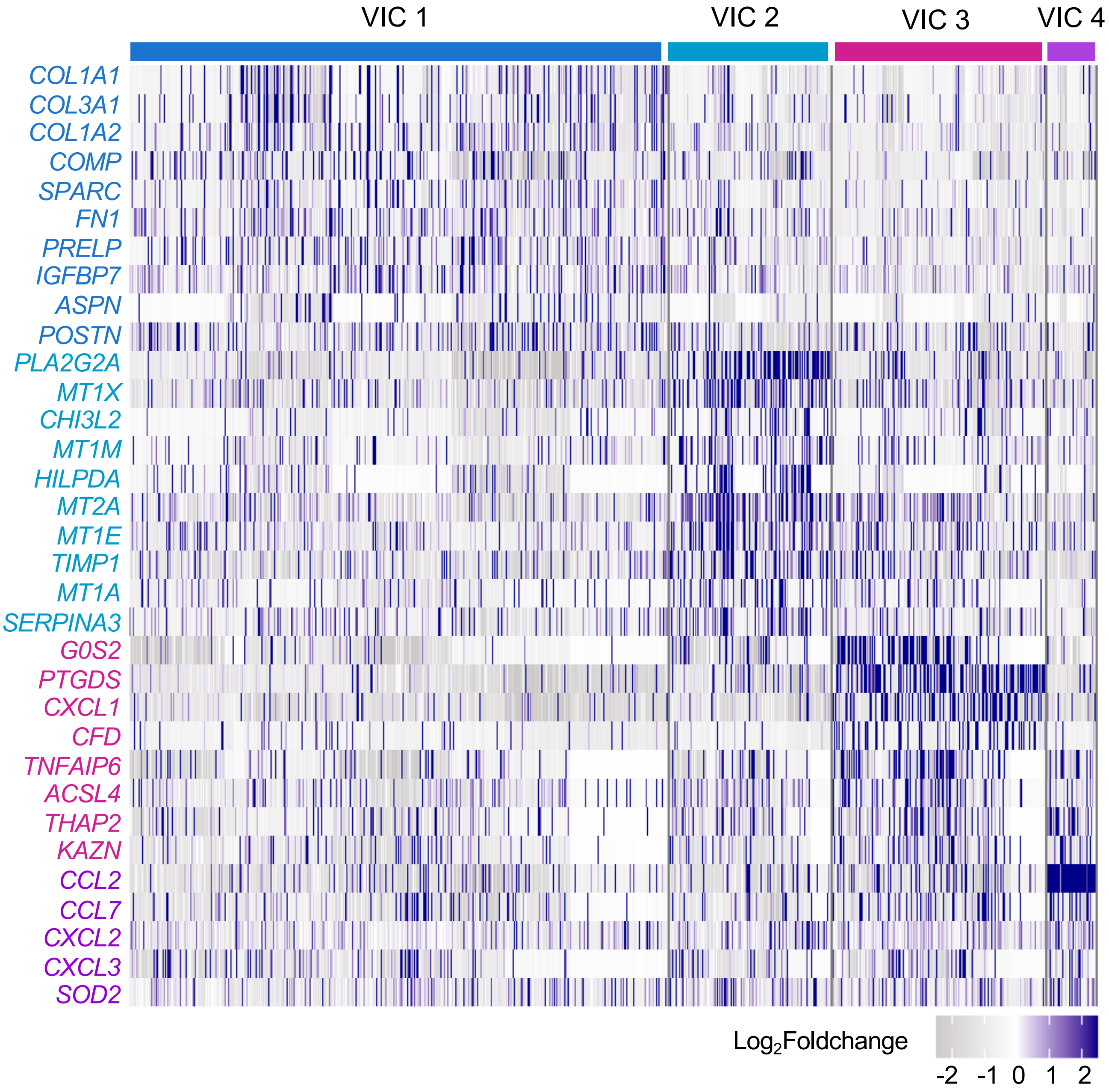


**Fig. VI: Heatmap of top 5 valvular interstitial cell markers.** Significance determined using Wilcoxon rank sum testing; $\log_2$Foldchange > |0.5|; *$P_{adj}$<0.0001.

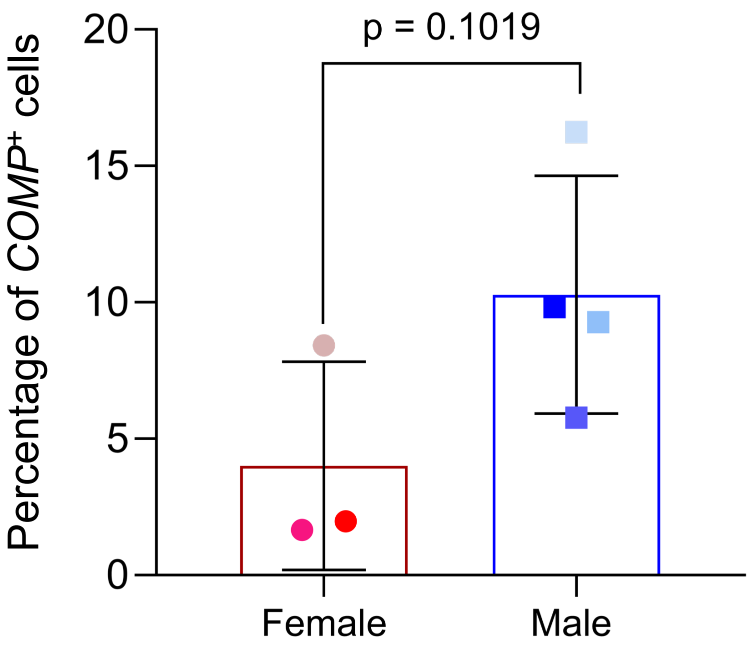


**Fig. VII: Percentage of *COMP*+ cells in male and female diseased tissue.** Sample size: N = 3 female diseased and N = 4 male diseased with at least 2 leaflets per patient. Significance tested with Welch's t-test (*$P < 0.05$). Mean ± S.D. shown.

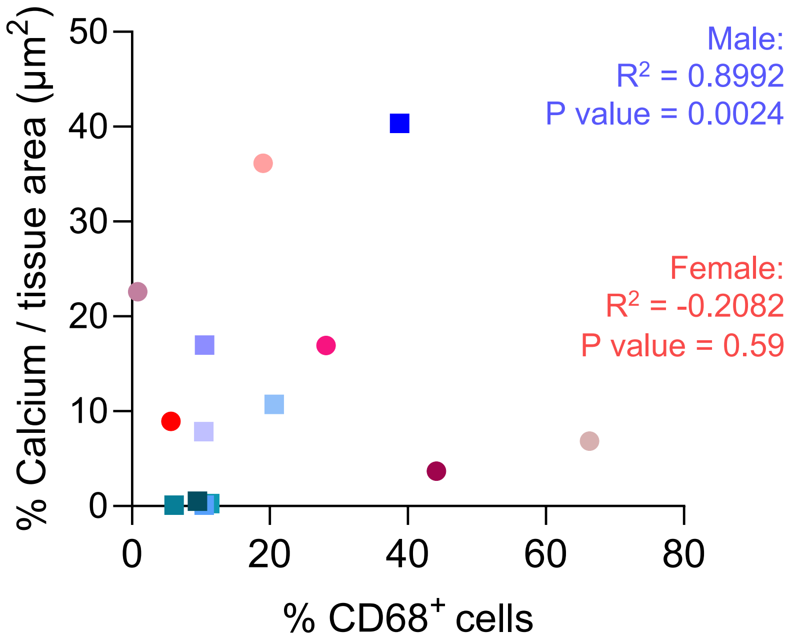


**Fig. VIII: Correlation of tissue calcification and $CD68^+$ cells in male and female tissue.** Sample size: N = 7 female tissues and N = 8 male tissues with at least 2 leaflets per patient. Correlation tested with Pearson's correlation coefficient.

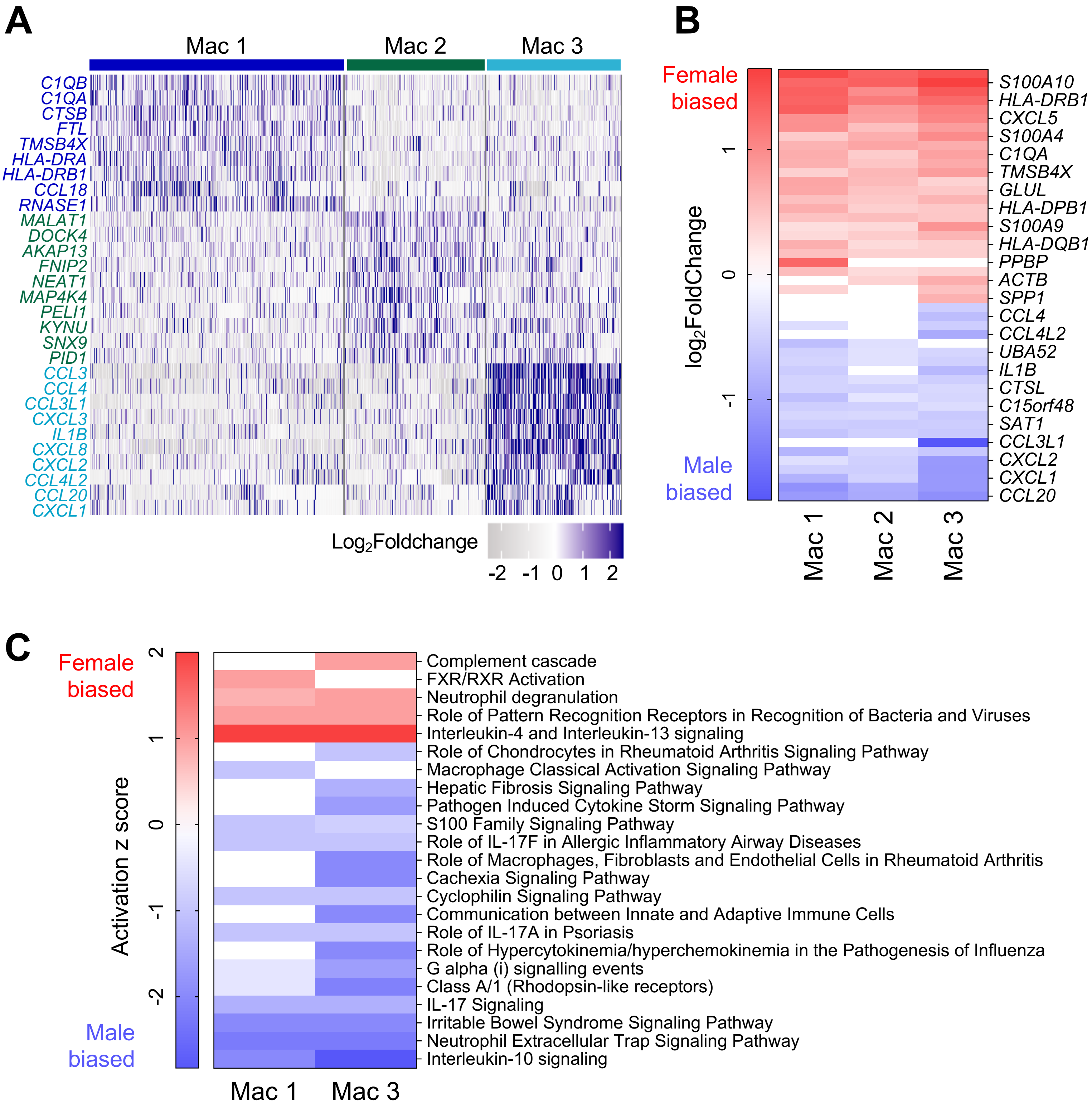


**Fig. IX: Macrophage subtype gene expression and associated pathways.** (A) Heatmap of top 10 markers per macrophage subtype. Significance determined using Wilcoxon rank sum testing; $\log_2$Foldchange > |0.5|; *$P_{adj}$<0.0001. (B) Differentially expressed genes between diseased male and diseased female macrophage subtypes. Significance determined using Wilcoxon rank sum testing; $\log_2$Foldchange > |0.5|; *$P_{adj}$<0.0001. (C) Ingenuity Pathway Analysis of pathways associated with mac 1 and mac 3 subtypes. ; *$P_{adj}$<0.05; activation z-score > |1| .

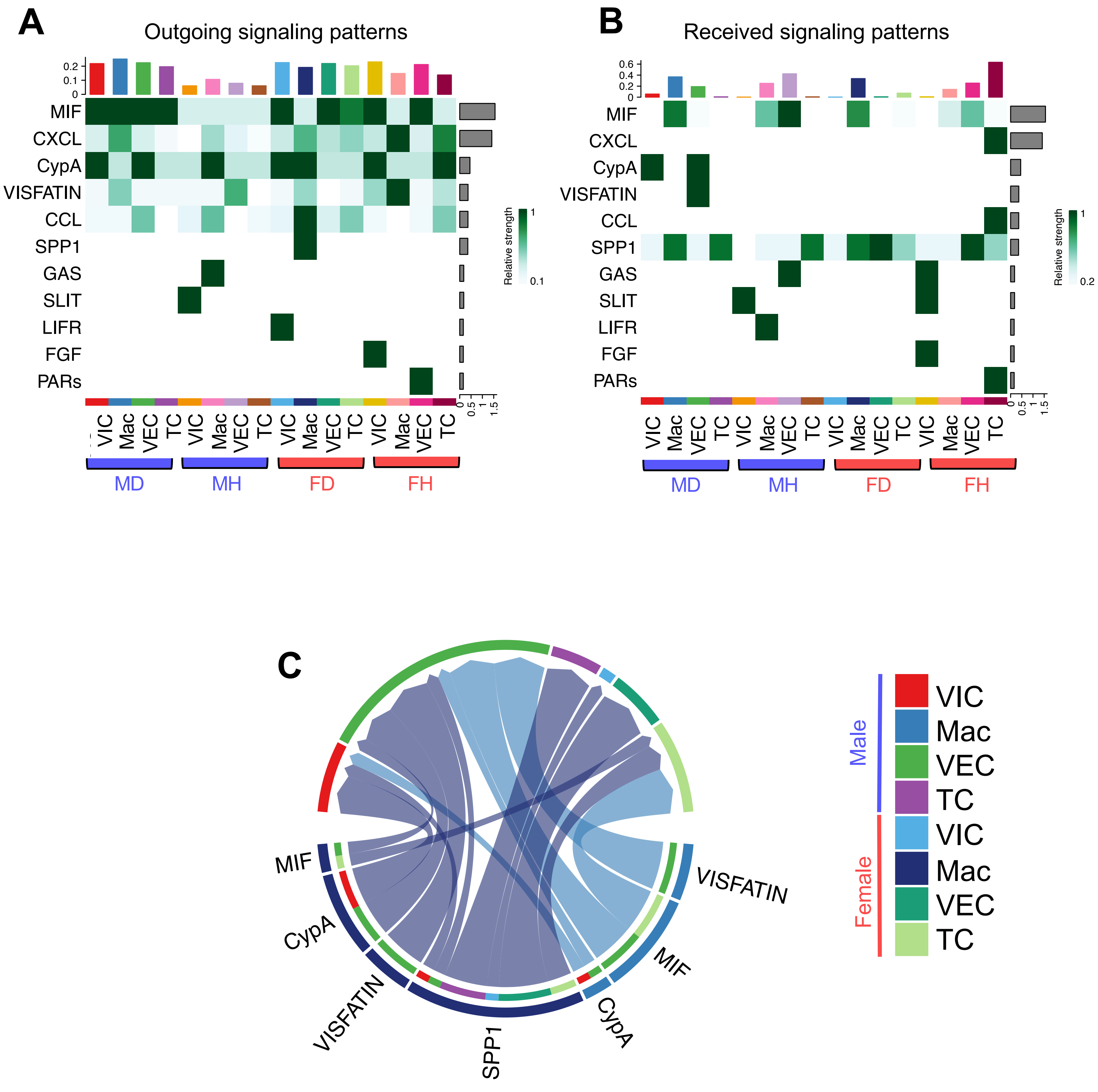


**Fig. X: Signaling patterns among single cell RNA sequencing clusters.** (A) Outgoing signaling patterns among all cell types. Sample size: N = 1 female healthy, N = 3 female diseased, N = 4 male healthy, N = 4 male diseased**.** (B) Received signaling patterns among all cell types. Sample size: N = 1 female healthy, N = 3 female diseased, N = 4 male healthy, N = 4 male diseased**.** (C) Chord diagram depicting all signaling patterns among cell types. Sample size: N = 1 female healthy, N = 3 female diseased, N = 4 male healthy, N = 4 male diseased.

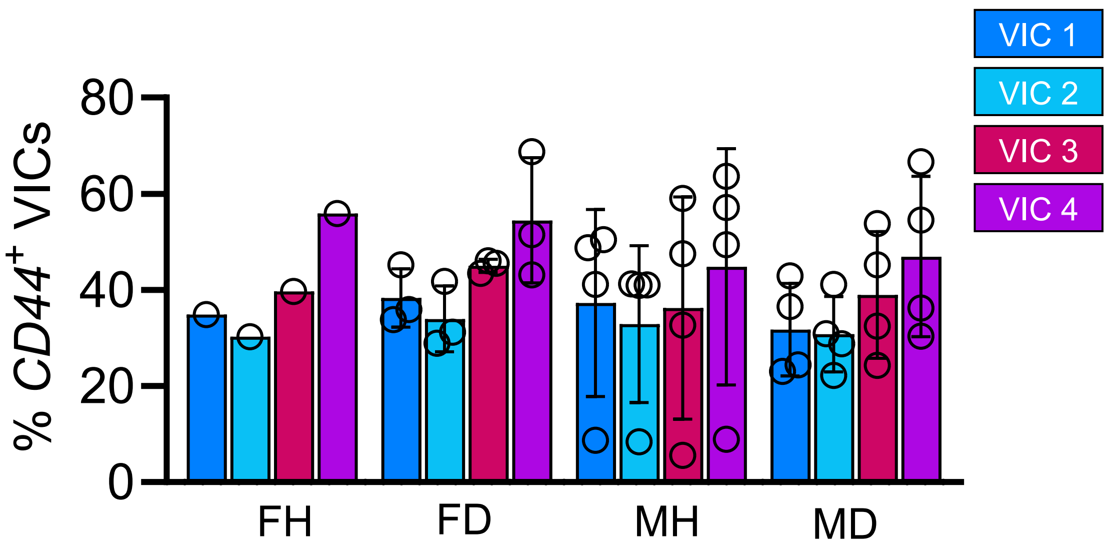


**Fig. XI: Percentage of *CD44* positive cells in VIC subtypes.** Sample size: N = 1 female healthy, N = 3 female diseased, N = 4 male healthy, N = 4 male diseased.

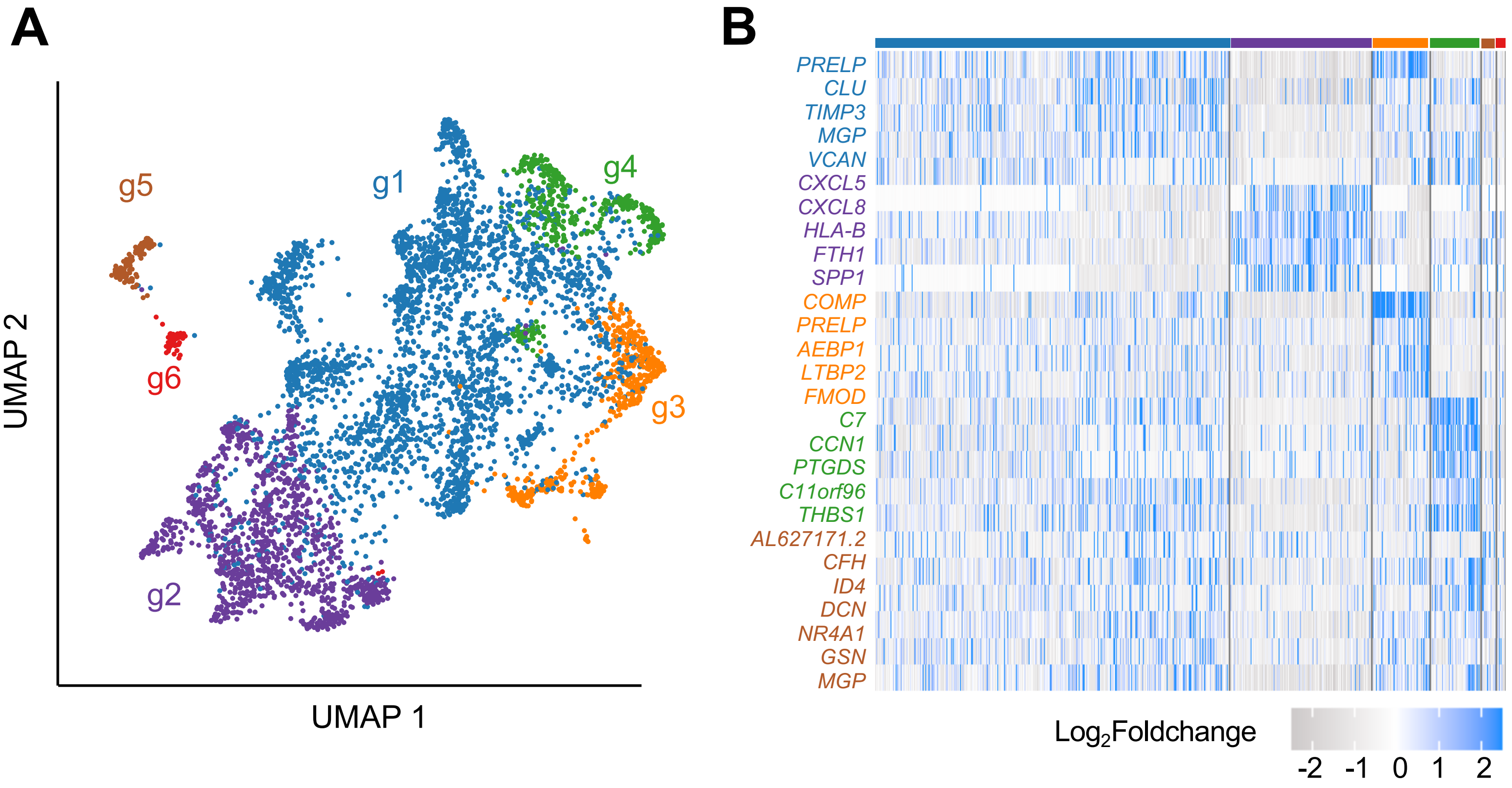


**Fig. XII: Unsupervised clustering of spatial transcriptomics RNA spots.** (A) UMAP of RNA spot clusters. (B) Heatmap of top 5 genes per RNA spot cluster. Significance determined using Wilcoxon rank sum testing; $\log_2$Foldchange > |0.5|; *$P_{adj}$<0.0001. Sample size: N = 1 male diseased and N = 1 female diseased with 2 tissue sections per biological replicate.

### Supplementary Table I. Patient clinical data

| PatientID | Sex | Status | Age | Diabetes | Hypertension | CAD | AVS | Atherosclerosis | BMI | Tabacco use |
|---|---|---|---|---|---|---|---|---|---|---|
| Patient 0 | Male | Healthy | 59 | | | | | | unk | unk |
| Patient 1 | Male | Diseased | 85 | | | | | | 20.7 | none |
| Patient 2 | Male | Healthy | 47 | | | | | | 24.91 | none |
| Patient 3 | Female | Diseased | 73 | • | • | | • | | 43.22 | none |
| Patient 4 | Male | Diseased | 79 | • | • | • | • | | 42.39 | none |
| Patient 5 | Male | Diseased | 76 | | | | | | 20.97 | none |
| Patient 6 | Female | Diseased | 76 | • | • | • | • | • | 25.74 | yes |
| Patient 7 | Male | Healthy | 60 | | | | | | 21.00 | none |
| Patient 8 | Male | Diseased | 65 | | | | | | 25.94 | yes |
| Patient 9 | Female | Diseased | 82 | • | • | | • | | 27.21 | yes |
| Patient 10 | Male | Healthy | 42 | | | | | | 20.31 | none |
| Patient 11 | Female | Healthy | 60 | | | | | | 19.48 | none |
| Patient 12 | Male | Diseased | 72 | | | | | | 23.92 | none |
| Patient 14 | Female | Healthy | 69 | | | | | | 24.98 | none |
| Patient 15 | Male | Diseased | 67 | | | | | | 26.15 | yes |
| Patient 18 | Female | Diseased | 75 | | • | | | | 18.04 | yes |
| Patient 19 | Male | Diseased | 80 | • | | | | | 26.08 | yes |
| Patient 20 | Male | Diseased | 80 | | • | • | | | 17.06 | yes |
| Patient 21 | Female | Diseased | 75 | • | • | • | | | 24.37 | none |
| Patient 22 | Male | Healthy | 60 | | | unk | unk | unk | unk | unk |
| Patient 23 | Female | Diseased | 63 | | | | | | 22.29 | yes |
| *Patient 24 | Male | Healthy | 50 | | | unk | unk | unk | unk | unk |
| *Patient 25 | Female | Healthy | 48 | | | unk | unk | unk | unk | unk |
| *Patient 26 | Female | Diseased | 52 | | | unk | unk | unk | unk | unk |
| *Patient 27 | Female | Diseased | 54 | | | unk | unk | unk | unk | unk |
| *Patient 28 | Male | Diseased | 50 | | | unk | unk | unk | unk | unk |

*Publicly available data from Xu et. al[22]

### Supplementary Table II. Sample sizes for all experimental procedures

| Patient ID | Sex | Status | Masson's Trichrome | Alizarin Red | SEM | scRNA seq | Spatial Transcriptomics | RNA-FISH | Protein staining |
|---|---|---|---|---|---|---|---|---|---|
| Patient 0 | Male | Healthy | • | • | | | | | |
| Patient 1 | Male | Diseased | • | • | • | • | • | • | • |
| Patient 2 | Male | Healthy | • | • | | • | | | • |
| Patient 3 | Female | Diseased | • | • | • | | | | • |
| Patient 4 | Male | Diseased | • | • | • | • | | | |
| Patient 5 | Male | Diseased | • | • | • | • | | | • |
| Patient 6 | Female | Diseased | • | • | • | • | • | • | • |
| Patient 7 | Female | Healthy | • | • | | | | | • |
| Patient 8 | Male | Diseased | • | • | | | | | • |
| Patient 9 | Female | Diseased | • | • | | | | | • |
| Patient 10 | Male | Healthy | • | • | | • | | | • |
| Patient 11 | Female | Healthy | • | • | • | | | | • |
| Patient 12 | Male | Diseased | • | • | | | | | |
| Patient 14 | Female | Healthy | • | • | | | | | • |
| Patient 15 | Male | Diseased | • | • | • | | | • | • |
| Patient 18 | Female | Diseased | • | • | • | | | • | • |
| Patient 19 | Male | Diseased | • | • | | | | • | • |
| Patient 20 | Male | Diseased | • | • | | | | • | • |
| Patient 21 | Female | Diseased | • | • | • | | | | • |
| Patient 22 | Male | Healthy | • | • | • | • | | • | • |
| Patient 23 | Female | Diseased | | | | | | | • |
| *Patient 24 | Male | Healthy | | | | • | | | |
| *Patient 25 | Female | Healthy | | | | • | | | |
| *Patient 26 | Female | Diseased | | | | • | | | |
| *Patient 27 | Female | Diseased | | | | • | | | |
| *Patient 28 | Male | Diseased | | | | • | | | |

### Supplementary Table III. Single cell RNA sequencing results statistics

| Patient ID | Number of reads | Mean reads per cell | Valid barcodes | Q30 bases in barcode | Q30 bases in RNA read | Q30 bases in UMI |
|---|---|---|---|---|---|---|
| Patient 1 | 171,820,731 | 406,196 | 91.8% | 95.8% | 93.1% | 97.0% |
| Patient 2 | 145,069,407 | 27,701 | 96.40% | 96.0% | 95.0% | 97.0% |
| Patient 4 | 272,696,391 | 38,697 | 96.6% | 95.8% | 94.2% | 97.0% |
| Patient 5 | 53,475,139 | 60,288 | 93.2% | 97.2% | 96.7 % | 98.1% |
| Patient 6 | 302,143,026 | 20,005 | 95.9% | 95.9% | 95.3% | 97.1% |
| Patient 10 | 132,404,712 | 131,615 | 96.1% | 95.9% | 93.0% | 97.10% |
| Patient 22 | 141,454,707 | 13,254 | 96.8% | 96.1% | 95.1% | 97.2% |

### Supplementary Table IV. Single cell RNA genomic mapping statistics

| Patient ID | Reads mapped confidently to transcriptome | Reads mapped confidently to exonic regions | Reads mapped confidently to intronic regions | Reads mapped confidently to intergenic regions | Sequencing saturation |
|---|---|---|---|---|---|
| Patient 1 | 61.7% | 57.1% | 8.1% | 5.1% | 93.9% |
| Patient 2 | 79.6% | 69.3% | 19.5% | 5.7% | 70.8% |
| Patient 4 | 82.1% | 75.7% | 12.7% | 5.4% | 84.3% |
| Patient 5 | 72.0% | 65.4% | 16.2% | 6.4% | 80.3% |
| Patient 6 | 81.6% | 74.1% | 13.9% | 5.0% | 76.2% |
| Patient 10 | 73.1% | 66.2% | 17.0% | 5.6% | 96.8% |
| Patient 22 | 75.2% | 63.8% | 26.8% | 3.9% | 83.7% |

### Supplementary Table V. Single cell RNA sequencing gene expression statistics

| Patient ID | Estimated number of cells | Fraction reads in cells | Mean reads per cell | Median genes per cell | Total genes detected | Median UMI counts per cell |
|---|---|---|---|---|---|---|
| Patient 1 | 423 | 76.7% | 406,196 | 2,632 | 21,702 | 8,651 |
| Patient 2 | 5,237 | 90.8% | 27,701 | 1,945 | 27,602 | 4,466 |
| Patient 4 | 7,047 | 89.8% | 38,697 | 1,519 | 27,577 | 3,298 |
| Patient 5 | 887 | 58.7% | 60,288 | 1,725 | 23,363 | 3,744 |
| Patient 6 | 15,103 | 88.7% | 20,005 | 1,134 | 29,717 | 2,096 |
| Patient 10 | 1,006 | 83.6% | 131,615 | 997 | 21,011 | 1,786 |
| Patient 22 | 10,673 | 85.9% | 13,254 | 774 | 26,964 | 1,136 |

Supplementary Table VI. Spatial transcriptomics sequencing results statistics

| Patient ID | Number of reads | Valid barcodes | Q30 bases in barcode | Q30 bases in RNA read | Q30 bases in UMI |
|---|---|---|---|---|---|
| Patient 1A | 190,026,709 | 97.5% | 94.9% | 94.7% | 96.6% |
| Patient 1B | 158,555,030 | 97.4% | 95.1% | 94.7% | 96.8% |
| Patient 6A | 163,156,910 | 97.1% | 94.9% | 94.6% | 96.6% |
| Patient 6B | 138,816,339 | 97.2% | 95.5% | 94.7% | 96.8% |

Supplementary Table VII. Spatial transcriptomics genomic mapping statistics

| Patient ID | Reads mapped confidently to transcriptome | Reads mapped confidently to exonic regions | Reads mapped confidently to intronic regions | Reads mapped confidently to intergenic regions | Sequencing saturation |
|---|---|---|---|---|---|
| Patient 1A | 47.3% | 49.3% | 18.8% | 5.8% | 98.5% |
| Patient 1B | 37.2% | 38.7% | 17.1% | 6.0% | 97.1% |
| Patient 6A | 41.5% | 42.8% | 10.9% | 6.2% | 97.0% |
| Patient 6B | 42.2% | 43.5% | 9.0% | 4.7% | 95.3% |

Supplementary Table VIII. Spatial transcriptomics genomic mapping statistics

| Patient ID | Number of spots under tissue | Fraction reads in spots under tissue | Mean reads under tissue per spot | Median genes per spot | Total genes detected | Median UMI counts per spot |
|---|---|---|---|---|---|---|
| Patient 1A | 1,573 | 76.5% | 81,218 | 317 | 15,338 | 452 |
| Patient 1B | 1,264 | 67.9% | 73,381 | 428 | 15,612 | 644 |
| Patient 6A | 1,658 | 72.9% | 61,818 | 306 | 16,304 | 413 |
| Patient 6B | 1,693 | 75.4% | 54,482 | 493 | 17,023 | 746 |

Supplementary Table IX. Sex differences in risk factors

| Variables | Male (N=12) | Female (N=9) | *P* value |
|---|---|---|---|
| Age, years | $63.69 \pm 14.71$ | $66.88 \pm 10.29$ | 0.60 |
| Body Mass Index, $kg/m^2$ | $24.49 \pm 6.63$ | $26.15 \pm 8.24$ | 0.73 |
| Coronary artery disease, n (%) | 2 (17%) | 2 (22%) | $> 0.99$ |
| Hypertension, n (%) | 2 (17%) | 4 (44%) | 0.12 |
| Diabetes, n (%) | 2 (17%) | 4 (44%) | 0.61 |
| Tobacco Use, n (%) | 4 (33%) | 4 (44%) | $> 0.99$ |
| Aortic stenosis, n (%) | 1 (8%) | 2 (22%) | 0.27 |

### Supplementary Table X. Transcriptomics versus proteomics

| **Healthy Male** | | **Diseased Male** | | **Healthy Female** | | **Diseased Female** | |
|---|---|---|---|---|---|---|---|
| **Fibrotic proteins** | **Calcific proteins** | **Fibrotic proteins** | **Calcific proteins** | **Fibrotic proteins** | **Calcific proteins** | **Fibrotic protiens** | **Calcific protiens** |
| DCN | LAMA2 | LGALS1 | B2M | CRYAB | HLA-A | ACTB | C1QB |
| PRELP | C7 | PLA2G2A | BASP1 | ATP1B3 | PTGDS | APOE | CST3 |
| MGP | FN1 | SERPINE2 | HLA-A | SNX9 | | LGALS1 | CTSB |
| ANXA1 | SEPINA3 | MIF | IGFBP4 | CYCS | | MGP | FN1 |
| TIMP3 | | TPI1 | MANF | NAMPT | | ANXA1 | HLA-DPB1 |
| HSPA1A | | PKM | MYL6 | AKR1C1 | | S100A6 | HLA-DRA |
| HSP90AA1 | | FTH1 | NDUFA4 | FILIP1L | | TAGLN2 | HLA-DRB1 |
| CD81 | | EIF5A | PTMA | | | S100A10 | IGFBP4 |
| PLXDC2 | | H3F3B | TIMP1 | | | HSPA1A | MYL6 |
| IGFBP5 | | EEF1G | TYMP | | | DSTN | PTMA |
| HNRNPH1 | | S100A11 | | | | PPIB | SPARC |
| | | HINT1 | | | | CRIP1 | TIMP1 |
| | | ATP1B3 | | | | PKM | |
| | | HLA-B | | | | FTH1 | |
| | | EIF4A1 | | | | CD9 | |
| | | CHMP4B | | | | COL3A1 | |
| | | CYCS | | | | CCL18 | |
| | | CD59 | | | | CSTB | |
| | | AKR1C1 | | | | HSPD1 | |
| | | SRP14 | | | | HSP90AA1 | |
| | | COX4I1 | | | | ARPC3 | |
| | | CD63 | | | | CD81 | |
| | | PDCD5 | | | | S100A11 | |
| | | | | | | S100A4 | |
| | | | | | | CLIC1 | |
| | | | | | | GNAS | |
| | | | | | | NACA | |
| | | | | | | CD63 | |
